# Robustness of biomolecular networks suggests functional modules far from the edge of chaos

**DOI:** 10.1101/2023.06.30.547297

**Authors:** Kyu Hyong Park, Felipe Xavier Costa, Luis M Rocha, Réka Albert, Jordan C Rozum

## Abstract

A common feature of complex systems is their ability to balance the flexibility needed to adapt to their environment with the rigidity required for robust function. It has been conjectured that living systems accomplish this by existing at the “edge of chaos”, i.e., the critical boundary between ordered and disordered dynamics. Simple toy models of gene regulatory networks lend support to this idea, and mathematical tools developed for these toy models yield similar results when applied to experimentally-supported models of specific cellular regulatory mechanisms (functional modules). Here, however, we demonstrate that a deeper inspection of 72 experimentally-supported discrete dynamical models of functional modules reveals previously unobserved order in these systems on long time scales, suggesting greater rigidity in these systems than was previously conjectured. Our analysis relies on new measures that quantify the tendency of perturbations to spread through a discrete dynamical system. A benefit of our new approach is that it accounts for how system trajectories are mapped to phenotypes in practice. Because these measures are computationally expensive to estimate, existing tools were insufficient for the ensemble of models considered here. To simulate the tens of millions of trajectories required for convergence, we developed a multipurpose CUDA-based simulation tool, which we have made available as the open-source Python library cubewalkers. We find that in experimentally-supported models of biomolecular functional modules, perturbation propagation is more transitory than previously thought, and that even in cases where large perturbation cascades persist, their phenotypic effects are often minimal. Moreover, by examining the impact of update scheme on experimentally-supported models, we find evidence that stochasticity and desynchronization can lead to increased recovery from regulatory perturbation cascades in functional modules and uncover previously unreported population-level robustness to even timing perturbations in these systems. We identify specific biological mechanisms underlying these dynamical behaviors and highlight them in experimentally-supported regulatory networks from the systems biology literature. Based on novel measures and simulations, our results suggest that–contrary to current theory–functional modules of biological systems are ordered and far from the edge of chaos.

## I. INTRODUCTION

“The edge of chaos,” a term coined by Packard in 1988 [44], refers to the tendency of adaptive systems to evolve toward a dynamical regime that lies between order and disorder. In systems biology, this is often referred to as the criticality hypothesis [56], and is closely related to work by Kauffman [32, 48] and Derrida [16, 17], who demonstrated that simple tunable models of gene regulation exhibit an order-to-chaos phase transition. Near this transition, it is conjectured, living systems optimally balance the rigidity required to function in a noisy environment with the flexibility required to undergo developmental, metabolic, and evolutionary processes that depend on cellular context. Dynamically, the boundary between order and disorder is often understood through the lens of trajectory separation; here, we seek to understand it through the lens of phenotypic fragility and its inverse counterpart, robustness.

The fragility of a cellular phenotype describes how easily it transitions to a different phenotype, and determines, for example, a cell’s ability to differentiate, its susceptibility to oncogenesis, and the fidelity of its signal processing.

This has been measured experimentally by genetically or pharmacologically perturbing genes and measuring the impact on cellular phenotypes [7, 25, 31]. In the context of dynamical models of biomolecular processes, the traditional approach to understanding phenotypic fragility is inspired by the analysis of random Boolean networks (RBNs) and considers the propagation of a large, temporary disruption to an individual component of the system (e.g., the depletion of a protein) [48]. In other words, an initial perturbation, on average, decays to extinction in the long-term dynamics of ordered (robust) systems, but it grows and spreads globally in the disordered (fragile) case. In RBNs, the average short-term propagation of initial perturbations, as measured by the Derrida coefficient, is sufficient to determine the average long-term spreading behavior [16, 17]. It therefore indicates the critical boundary between the ordered and disordered dynamical regimes in RBNs, which occurs when its value is 1 [51].

Non-random, experimentally-supported Boolean networks are popular tools for modeling biomolecular functional modules (regulatory mechanisms governing specific cellular processes) [2, 29]. As more of these models are constructed, one can ask whether an ensemble of such models exhibits properties similar to those of RBNs. In fact, many do have Derrida coefficients near 1 [14, 37, 55]. This observation lends support to the criticality hypothesis, but some caution is required; the connection between short-term perturbation response (Derrida coefficient) and the long-term perturbation explosion or extinction is not as straightforward in non-random models. In this work, we challenge the assertion that existing non-random Boolean models cluster on the boundary between order and disorder by using biologically-grounded measures of phenotypic fragility. Our analysis of these models reveals highly ordered perturbation responses that are obfuscated in the usual approach based on the Derrida coefficient and trajectory separation. We show that the criticality hypothesis is not valid in a battery of experimentally-supported models of biomolecular networks, which represent the state-of-the-art in causal modeling in systems biology (see below). Because these networks model subsystems of whole organisms studied in isolation, our results suggest that for the criticality hypothesis to be true, criticality of living systems must arise as a meso-scale phenomenon, through the coupling of (ordered) functional modules or even populations of organisms.

Our testbed for this study is a curated collection of 72 experimentally-supported, peer-reviewed Boolean network models of biomolecular functional modules found in the Cell Collective database [29], which represents the independent efforts of dozens of research groups. In all of these models, each included regulatory interaction is tagged with an experimental justification from the systems biology literature. Each node in these Boolean networks corresponds to a specific biomolecular entity (e.g., gene, protein, or cellular subprocess). These nodes each have two possible states at any given time step, which represent the activity or inactivity of the corresponding entity (e.g., transcription of a gene, phosphorylation of a protein, or initiation of a cellular process). The states of the nodes are governed by Boolean update functions, which convert the states of a node’s regulators into a binary output. Time is usually modeled as an implicit variable in these systems, and there are various methods for scheduling the update of variables. Though the steady states of the network are independent of update scheme, the oscillatory behavior of the system is not [19, 21, 49].

Indeed, the update scheme has a dramatic impact on the long-term dynamics of random networks along the order-to-chaos critical boundary [23, 26, 47]. In non-random models, however, rich dynamical behaviors can persist across update schemes, as illustrated in [15], though to our knowledge this has not previously been studied systematically. By thoroughly examining the impact of update scheme on experimentally-supported models, we characterize their response to perturbations in the timing and synchronization of regulatory events to explore population-level order and robustness in these systems.

In this work, we consider two extreme (and quite common) schemes: synchronous update and asynchronous update. These schemes have various trade-offs and either can be valid or invalid depending on modeling context. In the synchronous update, every node updates its state every time step. In other words, the state of each node at time *t* + 1 is determined by the state of its regulators at time *t*. This scheme produces fully deterministic dynamics. Due to various analytical and computational conveniences, synchronous update is a popular scheme for very large random models. Synchronous update treats all biomolecular events (e.g., gene transcription) as simultaneous, which can sometimes lead to spurious oscillations. A common approach to removing these oscillations is to consider asynchronous update schemes, though this risks destroying meaningful oscillations as well. Here, we consider a stochastic, asynchronous update scheme in which a single variable is randomly selected (uniformly) at each time step to be updated. This random selection introduces stochasticity into the dynamics and destabilizes delay-sensitive oscillations [21, 49]. Thus, the asynchronous update can be viewed as a kind of timing perturbation introduced to the synchronous update.

We also take special care in handling the effect of source nodes, which usually codify a cellular context or a signal external to the model. Though such nodes are common in the modeling literature, we demonstrate that they are statistically rare in random models. Moreover, we show that source nodes have a large impact on various measures of order in Boolean networks. From a dynamical perspective, a “temporary” perturbation to a source node is unique in that it will always become permanent; this stands in contrast to the behavior of constant nodes, which recover immediately after perturbation and are common in both random and experimentally-derived models. In many biological applications, a perturbation to a source node is fundamentally different than a perturbation within the core of the network because source nodes often summarize the collective activity of many external components.

We consider various measures of short-term and long-term perturbation spread in both synchronous and asynchronous update schemes and in the context of fixed or perturbable source nodes using simulations. Previous work has focused on the use of short-term perturbation dynamics and statistical arguments as an avenue to estimate long-term dynamics in large networks because of the immense computational burden of ensuring that long-term perturbation measurements converge [4, 16, 18, 61]. To meet this challenge and directly measure long-term perturbation growth in non-random models, we developed cubewalkers, a highly-parallel GPU-based simulation toolkit, allowing us to quickly simulate many thousands of trajectories in a network simultaneously. Our software innovations, combined with the dramatic improvements in computational power over the last several decades enable, for the first time, high-fidelity measurements of long-term perturbation dynamics in real-world Boolean networks with hundreds of nodes or more. These measurements are fundamental to demonstrating the true dynamical regime of experimentally-supported biomolecular networks.

## II. METHODS

### A. Boolean network dynamics at the individual and population level

Boolean networks describe the regulatory dynamics of each node *X* as a Boolean update function *X*^⋆^ denoting the state of *X* at a subsequent time step. We define two special types of node that have unique effects on the dynamics: constant nodes, which have update functions of the form *X*^⋆^ = 0 or *X*^⋆^ = 1, and source nodes, which have update functions of the form *X*^⋆^ = *X*. More generally, update functions utilize the logical operations “AND”, “OR”, and “NOT” which we denote by ∧, ∨, and ¬, respectively. Each Boolean system with *N* nodes induces a state transition graph whose 2^*N*^ nodes represent all possible system states and whose directed edges indicate that the parent state can be updated in one time step to attain the child (successor) state. The attractors of a Boolean system are the terminal strongly connected components of the state transition graph (i.e., they have no edges that exit the component). Point attractors (also called steady states) consist of a single state, and oscillatory attractors (also called complex attractors) contain more than one state. The simplest type of oscillatory attractor is a limit cycle, in which the system revisits states in a deterministic order. The states that can reach an attractor via edges or paths in the state transition graph make up the basin of attraction of the attractor. In each network, the set of possible attractors can strongly depend on the update scheme used. Indeed, one of the most fundamental biomolecular circuit motifs, mutual inhibition, exhibits such behavior. Consider two mutually inhibiting genes, *A* and *B* described by the simple Boolean network with update functions

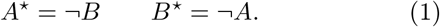

In the asynchronous update scheme, there are only two attractors: the steady states (*A, B*) = (1, 0) and (*A, B*) = (0, 1). In the synchronous update scheme, however, there is an additional oscillatory attractor that cycles between the states (*A, B*) = (0, 0) and (*A, B*) = (1, 1). Thus, the behavior of an individual instance of a model (i.e., a single cell) is highly sensitive to the timing of node update. This example highlights, however, that the *average* behavior of many instances (i.e., the population-level behavior) can be robust to update timing even when individual instances (cells) are not. To see this, consider the average activation value of gene *A* (by symmetry, the same analysis applies identically to gene *B*). Assuming uniformly sampled initial conditions and allowing enough time for convergence into an attractor, we observe that in the asynchronous scheme, an individual cell has a 50% probability of being in the (*A, B*) = (1, 0) steady state and a 50% probability of being in the (*A, B*) = (0, 1) steady state; thus, overall, the average value of *A* in the ensemble is 0.5. In the case of synchronous update, the system has a 25% probability of being in either steady state, and 50% probability of being in the oscillatory attractor. The average value of *A* (and also of *B*) in the oscillatory attractor, however, is 0.5, and thus, overall, the average value of *A* in the synchronous update is also 0.5, just as it is in the asynchronous case. This behavior need not hold in general. To quantify the extent to which this behavior occurs in the test models considered, we compare the converged average node values under synchronous and asynchronous update schemes and compute the root mean squared (RMS) difference between the synchronous and asynchronous average node values across all nodes of a model, which we discuss in detail in Section III A 2.

### B. Models considered

Throughout this work, we consider 72 models from the Cell Collective [29] and their dynamical properties. In some cases, nodes whose update functions are constants in the originally published version of a model have been reinterpreted as source nodes in the Cell Collective, or multiple source nodes have been merged. In such cases, we defer to the original publication; in most cases, this results in replacing the update functions for several source nodes with constant-value update functions. In addition, we correct a few typographical errors in the models, remove isolated nodes, and enforce constraints that were not previously enforced when multiple nodes encode more than two values of a single entity (e.g., low, medium, or high concentration of a protein). In all, 18 models are affected in some way. We use these modified versions of the models here in an attempt to more accurately capture the biology represented in these models. Overall, we observe very little difference in the distributions of the measures considered when compared to the unaltered Cell Collective ensemble, though for some measures, the differences in individual models can be large for measures that emphasize the role of source nodes (comparisons provided in Figure F.1 in Appendix F).

We also highlight several models with particularly interesting dynamical features. Throughout this work, these highlighted models are indicated by colored symbols. The shape of the symbols in various plots (whether highlighted or not) describes the biological category of the model whose parameters are plotted. This correspondence is summarized in Figure 1.

**FIG. 1.**
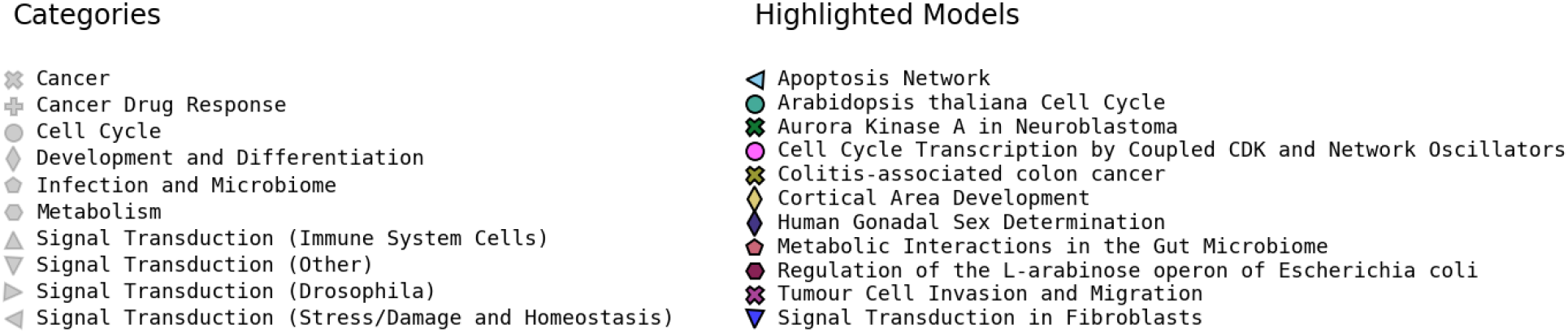
Legend indicating model categories (marker shape) and specific highlighted models (marker color).

### C. Simulation and analysis software

To compute various dynamical measures, including those introduced here, we developed the CUBEWALKERS Python library, a CUDA-based Boolean network simulator. It supports various update schemes (including user-specified schemes), node and edge control interventions, and probabilistic (e.g., noisy) update rules. To simulate Boolean networks, CUBEWALKERS parses Boolean update functions given either in algebraic form (as a string) or in the form of a lookup table (as a Boolean vector for the outputs and list of regulators). The parsed Boolean update rules are used to generate source code for a CUDA kernel that is compiled via the Python interface CUPY [41]. When simulating the dynamical model, CUBEWALKERS executes this kernel on an array of state vectors, with each state vector representing the values of the nodes in a single network simulation instance, or “walker”. Updates for the nodes of each walker are computed in parallel on the GPU for each time step according to the chosen update scheme. Various options for conserving memory are implemented, such as retaining only the most recent time points or tracking only averages across walkers.

Conducting unbiased quantitative benchmarks that compare the performance of CUBEWALKERS to that of other Boolean simulation tools is complicated by the fact that CUBEWALKERS is primarily GPU-based, while competing tools run entirely on the CPU. Therefore, one must assess the relative quality of the CPU and GPU used in benchmarking comparisons. Furthermore, most CPU-based Boolean simulation tools do not execute operations in parallel; however, user-side parallelization is often possible. Despite these caveats, the performance advantage of CUBEWALKERS is dramatic and convincing in practice. We compared against the Python library CANA (which wraps a C implementation via Cython) [11] and the Python library BOOLEANNET(which is written in pure Python) [3]. Detailed benchmark results are provided in Appendix A, and are summarized here. For synchronous simulations of models in the Cell Collective on consumer hardware, we demonstrate a speedup of approximately 350 times on average compared to simulation using CANA and a speedup of approximately 11, 000 times on average compared to simulation using BOOLEANNET. We also compared CUBEWALKERS performance to the performance of parallelized simulations using CANA and BOOLEANNET on a high-performance computing workstation. In this case, CUBEWALKERS outperforms CANA by a factor of 16.6 and outperforms BOOLEANNET by a factor of just under 550. Furthermore, CUBEWALKERS has approximately 10 times better performance on our consumer test hardware than is achieved using parallel simulations with CANA on our specialty high-performance hardware.

Note that the simulation results presented herein required several days of computer time using CUBEWALKERS, so the approximately 16-times slowdown we expect from the second-fastest software considered would result in months of excess computation.

We determine the number *W* of walkers (independent simulations) to use in numerical experiments by examining the convergence of average node values (i.e., the output value of each individual node averaged across the *W* independent simulations). At any given time point, the value of an individual node for an individual walker can be viewed as a Bernoulli random variable with an unknown success probability. While the values of two different nodes for the same walker are not independent in general, the values of the same node for two different walkers are always independent. Thus, the central limit theorem applies to the average node values: the standard deviation of each node’s average value is bounded by 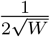 in the maximum variance case (attained for Bernoulli probability 0.5). In most experiments, we use at least *W* = 2, 500 independent simulations to obtain an expected standard deviation in the average node values of less than 0.01. This convergence is remarkable because it reveals that average node values can be accurately calculated in large network models using a relatively small sample size. In a network with 50 nodes, for example, a sample of *W* = 2, 500 initial states represents just over two trillionths of the state space, but is sufficient to calculate average node values at a given time step to within a few percent. Other measures we compute require more walkers to achieve the same desired accuracy; in the most extreme case, we used *W* = 800, 000 walkers. To determine the number of time steps to use in our simulations, we consider the agreement between four sub-intervals of a time-averaging window [*T*_*b*_, *T*_*b*_ + *T*_*w*_] for various choices of *T*_*b*_ and *T*_*w*_. We chose the number of time steps to simulate, *T* = *T*_*b*_ + *T*_*w*_, such that the largest per-node disagreement across sub-intervals was acceptably low for all Cell Collective models (below 0.0066 in the worst case, and significantly lower in most cases). Setting *T*_*b*_ = 50*N* +1 000 and *T*_*w*_ = 5*N* +5 000 is sufficient for computing the average behaviors of nodes in all but three models in the Cell Collective, which require longer simulation time. Further details and numerical tests supporting the mathematical arguments of this section are provided in Appendix B.

### D. Dynamical measures

The growth of small perturbations in Boolean networks is widely viewed as the hallmark of chaos in these systems [18]. In random models, this is often studied using the Derrida map, which relates the size of a perturbation at time *t*_0_ to the size of a perturbation at time *t*_0_ + 1. The Derrida map can be computed by sampling many pairs of initial states that differ in *h* variable values and evolving each pair of states using one synchronous time step. The average separation (Hamming distance) of the pairs becomes the numerical estimate for the value of the Derrida map at *h* [16, 17]. In principle, the states reached after one time step might not be distributed uniformly in the state space, so the Derrida map does not necessarily predict whether small perturbations grow or shrink in the long term. In random networks in the thermodynamic limit (*N* → ∞), however, whether the fixed point of the Derrida map is a finite fraction of the network is determined by the value of the map at *h* = 1. This value is called the *Derrida coefficient* and is equal to the average sensitivity of the network [14, 51]. Perturbations tend to spread to a finite fraction of the network only if the Derrida coefficient is greater than one; this corresponds to the chaotic regime. When the Derrida coefficient is less than one, the system is in the ordered regime in which perturbations tend to die out. A phase transition occurs on the critical boundary where the Derrida coefficient is equal to one. Dynamically, the Derrida coefficient can be defined as

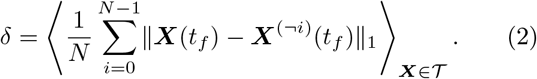

In this formula, ***X*** is a time-dependent vector of node states, **𝒯** is the set of all trajectories in the system, and ⟨· ⟩_***X****∈* ***𝒯***_ denotes the average taken over all possible trajectories, where the initial conditions and update schedules are sampled uniformly. The trajectory ***X***^(*¬i*)^(*t*)) is the trajectory that initially differs from ***X***(*t*) only in position *i* and is updated in the same way as ***X***(*t*) at every time step (this is important in stochastic update schemes). The comparison time, *t*_*f*_ is chosen such that *N* node updates are performed, and thus is equal to 1 in the synchronous update and to *N* in the asynchronous update. The summand ∥***X***(*t*_*f*_) − ***X***^(*¬i*)^(*t*_*f*_)∥_1_ is the L1-norm (absolute difference summed, or, for Boolean inputs, the Hamming distance) between ***X***^(*¬i*)^(*t*_*f*_)) and *X*(*t*_*f*_) at time *t*_*f*_.

In addition to the Derrida coefficient, *δ*, we consider three other measures to describe the response of systems to small (single-node) perturbations: final (average) Hamming distance *h*^∞^, quasicoherence *q*, and fragility *φ*. We illustrate the intuitive meaning of these measures in the case of a single-node oscillator *A*^⋆^ = ¬*A* in Figure 2.

The *final Hamming distance h*^∞^ is a direct measure of the long-term separation between trajectories that initially differ in a single node’s value. It is defined as

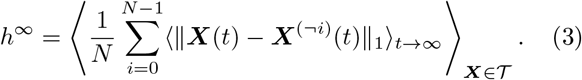

Here, ⟨·⟩_t→∞_ indicates the average taken from any finite initial time *t* = *t*_0_ to *t* = ∞; note that the value of the time average does not depend on the value of *t*_0_. Intuitively, *h*^∞^ measures the asymptotic separation (on average) between all trajectory pairs that initially differ in only one node value. Note that the Hamming distance

**‖*X***(*t*) ***X***^(*¬i*)^(*t*)‖_1_ does not necessarily converge for large *t* (it may oscillate), necessitating the time average calculation.

The *h*^∞^ measure is sensitive to phase shifts; if ***X***(*t*) and ***X***^(*¬i*)^(*t*)) converge to the same limit cycle, for example, but are offset, ***X***(*t*)‖***X***^(*¬i*)^(*t*)‖_1_ can be nonzero for all time even though the trajectories have the same long-term behavior. To distinguish this case from the case when ***X***(*t*) and ***X***^(*¬i*)^(*t*)) converge to different attractors, we propose two additional measures.

The first of these is the *fragility φ*, which we define as

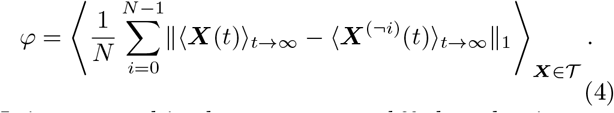

It is expressed in the same way as *h*^∞^, but the time averaging occurs inside the L1-norm, rather than outside it. This removes sensitivity to phase shift, and can be interpreted as a measure of separation in average values, rather than as an average separation. From a biological standpoint, this is desirable when a pair of trajectories with a high average separation but the same average behavior (as happens if the trajectories are time-shifted but otherwise identical) should be interpreted as phenotypically equivalent. Such trajectories may represent cells that are at different points of otherwise identical cell cycles. As a simple example, consider the system *A*^⋆^ =¬ *A*; *B*^⋆^ = *B*. Here, there are only two attractors in either update scheme: *A* will always oscillate, and *B* can be fixed in either value. If *B* is perturbed, the original and perturbed trajectories will always agree in *A* and differ in *B*, while if *A* is perturbed, the opposite is true and the system simplifies to the example of Figure 2. This conclusion holds in both synchronous and asynchronous update schemes because, in the latter, we constrain the selection of the update node to always be the same in both trajectories. Thus, *h*^∞^ = 1 for this system in both update schemes. In the case when *A* is perturbed, however, the average value of *A* does not differ between the two trajectories, and thus, as in the case of Figure 2, this perturbation contributes 0 to *φ*. As perturbations to *B* do alter the average value of *B*, they contribute 1 to *φ* and we therefore find *φ* = 0.5 in this system overall. This indicates that half of the long-term trajectory separation due to single-node perturbations stems from time-lag effects, which are not necessarily biologically relevant.

**FIG. 2.**
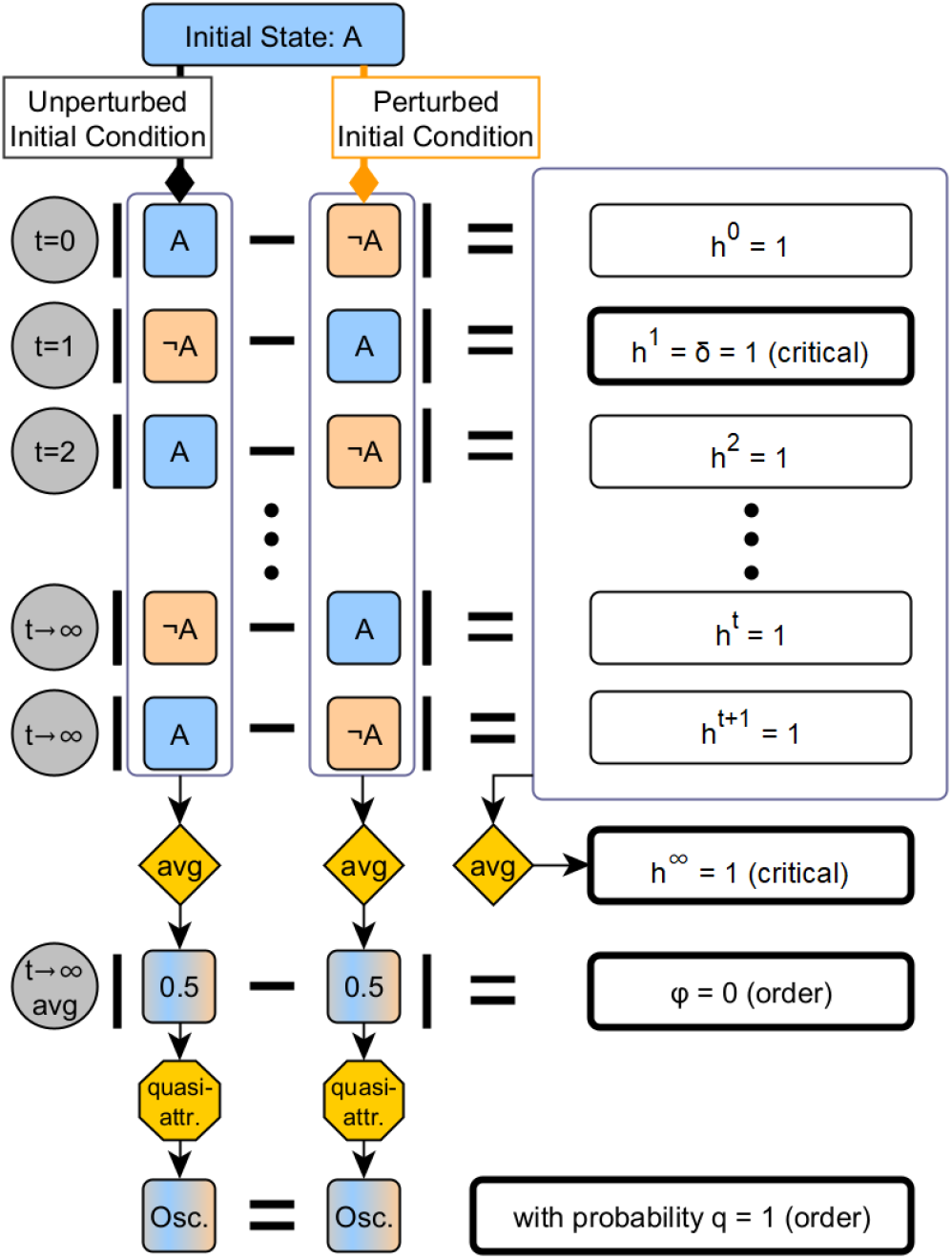
Comparison of four perturbation response measures for a one-node oscillator. The unperturbed oscillator alternates between two states: its initial state *A*, which could be 0 or 1, and the opposite state, ¬*A*, which is 1 if the initial state is 0, and 0 if the initial state is 1. The perturbed trajectory begins with the oscillating node in the opposite state compared to the unperturbed trajectory, but otherwise its time evolution proceeds in the same fashion. At each time step *t*, the Hamming distance *h*^*t*^ is computed. In the special case of *t* = 1, *h*^1^ is the Derrida coefficient *δ*, which evaluates to 1 in this case. Indeed, *h*^*t*^ = 1 for all *t*, so the asymptotic average of the Hamming distance, which we call the final Hamming distance (denoted *h*^∞^) evaluates to 1 as well. Alternatively, we can compute and compare the average behavior of the two trajectories. In both cases, the node is in the 0 state for half of the time steps, and in the 1 state in the other half. Thus, the average node value is 0.5 for both trajectories. Thus the fragility *φ*, defined as the difference in these averages, is 0. Furthermore, we can consider a more coarse-grained averaging, where we only record the probability that a randomly perturbed node (in this example there is only one node to choose from) results in a different quasiattractor, i.e., a different pattern of fixed and oscillating nodes. This probability is called quasicoherence. In this case, perturbing the initial state always results in the same quasiattractor (in which the sole node oscillates), so the quasicoherence is 1.

Another measure that can distinguish phenotypic differences from phase shifts is the *quasicoherence q*, which is closely related to the coherence measure introduced in [59]. Coherence is defined as the fraction of (***X***(*t*), ***X***^(*¬i*)^(*t*)) pairs that converge to the same attractor; in [59], coherence was defined only for synchronous update, but the extension to the asynchronous case is trivial. The primary barrier to adopting coherence as a measure is that attractor identification can be computationally expensive; sometimes prohibitively so. We therefore define and adopt quasicoherence as an alternative, which is defined as the fraction of (***X***(*t*), ***X***^(*¬i*)^(*t*)) pairs that converge to the same quasiattractor. Slightly modifying the convention of [60], we define a quasiattractor to be a pattern of fixed node values and oscillating nodes exhibited by an attractor. Two (or more) attractors may correspond to the same quasiattractor if they share the same set of active nodes, the same set of inactive nodes, and the same set of oscillating nodes. As a simple example, consider *A*^⋆^ = *B*; *B*^⋆^ = *C*; *C*^⋆^ = *A*. In the synchronous update, this system has four attractors: {000}, {111}, {001, 010, 100}, and {110, 101, 011}. In contrast, there are only three quasiattractors: 000, 111, and *⋆ ⋆ ⋆*, where *⋆* denotes that the node oscillates in all attractors that correspond to the quasiattractor. The quasicoherence can be written as

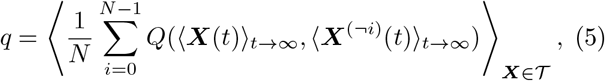

where *Q* : [0, 1]^*N*^× [0, 1]^*N*^ →0, 1 is defined such that *Q*(***X, Y***) is one if for all indices *i*, it holds that ***X***_*i*_ = 1⇔***Y*** _*i*_ = 1 and ***X***_*i*_ = 0⇔***Y*** _*i*_ = 0; otherwise *Q*(***X, Y***) is zero. The quasicoherence is 1 if all perturbed trajectories converge to the same quasiattractor as their unperturbed counterparts, and it is 0 if an initial perturbation to a single node always results in a different quasiattractor.

The quasicoherence, unlike the final Hamming distance and fragility, does not distinguish between the case when trajectories converge to very similar (but not equal) steady states from the case when they converge to very different steady states. Because the time averaging is conducted before comparison, it is not sensitive to phase shifts either. The quasicoherence is useful when long-term changes in the expression of even a small number of genes are phenotypically important. The fragility and quasicoherence are related to each other in that the fragility can be interpreted as a rescaled “fuzzy” version of the quasicoherence, as explained in Appendix C.

We compute these dynamical measures (*h*^∞^, *q*, and *φ*) numerically for each network in the Cell Collective using a simulation-based approach. First, we sample 2 500*N* initial states, produce a copy of each, and perturb each copy in exactly one node (for a total of 5 000*N* initial states). Each initial state is evolved forward in time for *T* = *T*_*b*_ + *T*_*w*_ time steps, and the various time averages are taken over the last *T*_*w*_ time steps, as described in section II C and in further detail in the Appendix B. This is done in both the synchronous and asynchronous update schemes. The Derrida coefficient is computed using one synchronous time step or *N* asynchronous time steps using 100 000 initial samples (for a total of 200 000 initial states when considering the perturbation).

In addition, to probe the effect of source nodes (nodes whose update functions are of the form *A*^⋆^ = *A*) in Boolean networks, we consider “fixed source” versions of these five measures in which the perturbed nodes may not be source nodes and in which all instances of *N* in the formulae are replaced by the number of nodes that are not source nodes. Importantly, constant nodes remain perturbable in these cases, as do nodes that become fixed as a direct consequence of the source node values. All other parameters are unchanged.

Taken together, this results in four variations of each measure: two possible choices of update, indicated by a subscript *s* for synchronous and *a* for asynchronous, and two possible choices for how to treat source nodes, indicated by subscript *f* or *p* for fixed source nodes or perturbable source nodes, respectively. For example, *φ*_*s,f*_ indicates the fragility computed using the synchronous update and not allowing for source nodes to be perturbed, while *φ*_*a,p*_ indicates the fragility computed using the asynchronous update and allowing source nodes to be perturbed. In total, we consider sixteen measures of node perturbation response. The four variants of the Derrida coefficient *δ* measure short-term perturbation response. The four variants of the final Hamming distance *h*^∞^ measure long-term perturbation response in a manner that is sensitive to phase shifts. The four variants of the fragility *φ* measure long-term perturbation response in a manner that is insensitive to phase shifts. Finally, the four variants of the quasicoherence *q* measure the probability that a node perturbation induces a long-term change in quasiattractor.

## III. RESULTS

### A. The effects of synchronization perturbation

We first consider the effects of perturbations to the synchrony of biomolecular events. By comparing network dynamics under synchronous and asynchronous update, we consider an extreme version of this timing perturbation in which no two nodes states can update simultaneously. We study this at the level of single networks (akin to studying individual cells) and at the level of network populations (akin to studying populations of cells). At the level of individual networks, we examine the effect of perturbations on the range of possible long-term behaviors, whereby a reduction of this range corresponds to increased order. At the population level, a synchronously updated network is timing robust if it retains the average population-level behavior even when the synchrony of the biomolecular events it encodes is disrupted. In other words, a Boolean network exhibits a robust and ordered response to timing perturbations at the population level if its average node values do not depend (much) on the choice of update scheme.

#### 1. Synchrony perturbation confers order by destroying attractors

The attractor repertoire of Boolean models (and specifically, the oscillatory attractors) depend on the update scheme [19, 49]. In general, there are more attractors under synchronous update than under asynchronous update. As synchronous update is deterministic, its oscillatory attractors are always limit cycles. Attractors that only exist for synchronous update rely on the exact timing of updates (such that multiple nodes change state at the same time), and disappear in case of variations of the update timing, causing the system to have more orderly behavior [21]. We identify several models in the Cell Collective with this property and characterize the mechanisms underlying it by studying simplified models that are obtained by percolating the fixed value of source nodes, on eliminating a self-edge-free node and plugging in its update function into the function of its targets [40, 57], and on merging nodes with similar regulatory roles. In this section, we highlight some of these models and describe the update dependence of their attractor dynamics. In some cases, these attractors can have a significant biological meaning, for example, they may represent the cell cycle; in other cases, these attractors are spurious. Readers interested in ensemble properties only may safely skip ahead to Section III A 2.

The Cell Cycle Transcription by Coupled CDK and Network Oscillators 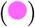 model [42] incorporates the known interactions among 9 cell cycle transcription factors and is one of several variants studied by Orlando et al. In synchronous update, this model has a point attractor corresponding to the G0 checkpoint and an oscillatory attractor that reproduces the sequence of transcription during the phases of the cell cycle. We find that the oscillatory attractor disappers under asynchronous update. To better understand the mechanisms that lead to this timing perturbation sensitivity, we simplified the model by merging closely related nodes and verified that the simplified model reproduced the correct transcription sequence under synchronous update. A key feature of the resulting network is that it consists of a positive feedback loop that intersects a shorter negative feedback loop. In general, this property ensures that the system is not monostable under synchronous update [5], i.e., an attractor other than the G0 fixed point exists. This extra attractor relies on synchrony and is therefore not robust to timing perturbation. The simplest example with these features is given in Equation 6; adding a delay node (Equation 7) equalizes the feedback loop lengths and results in monostability under synchronous update, consistent with the results of [5]. In asynchronous update, both systems are monostable. See Figure F.2 in Appendix F for further details.

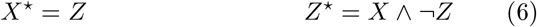

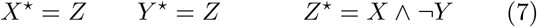

The Aurora Kinase A in Neuroblastoma 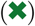 model developed by Dahlhaus et al. [13] exhibits a similar phenomenon. In that case, an attractor corresponding to defective mitosis (which leads to mitotic catastrophe and ultimately results in cell death) is present in synchronous update, but vanishes under timing perturbations. In a simplified model, we observe that the defective mitosis attractor requires a delay between the production and activation of the Aurora Kinase A protein. Notably, this attractor plays an important clinical role in neuroblastoma prognosis where defective mitosis is desirable. See Appendix D for a more detailed description of the biological and dynamical properties of this model, and for a simplified model that captures key properties of the system.

In contrast to the previous examples, the Regulation of the L-arabinose operon in *Escherichia coli* 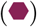 model [1] has spurious synchronous attractors that disappear under asynchronous update and do not have biological meaning. This model describes the regulation of the genes involved in arabinose metabolism in *E. coli*. In the input configuration corresponding to a medium level of external arabinose, available unbound AraC protein, and no external glucose, there are two point attractors, and four additional cyclic attractors under synchronous update. As in the example of Equation 1, these additional synchronous attractors arise from a positive feedback loop (here formed by four nodes). The original article describes these additional attractors as artifacts of the synchronous update, in contrast to the two biologically justified point attractors shared by both updates.

These models illustrate that the biological interpretation of a Boolean network can depend strongly on update scheme. Timing perturbations can destabilize oscillations that depend on specific delays between events by making them stochastic. This can lead to a decrease in the range of behaviors available to individual cells, ultimately resulting in dynamics that are more constrained and orderly.

#### 2. Timing-robust order emerges in cell populations

Though the attractor repertoire of models can be sensitive to update scheme at the level of individual cells, we observe that robustness to timing perturbations typically emerges at the cell population level. This suggests that populations of cells exhibit order that is not necessarily observable at the individual level. In almost all cases, the difference between the converged average node values in the synchronous and asynchronous updates is extremely small (see Figure 3). Notable exceptions include the Colitis-associated Colon Cancer 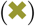, Aurora Kinase A in Neuroblastoma 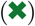, and Cortical Area Development 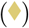 models. These three models have the three highest values of RMS difference and thus exhibit the least orderly response to timing perturbation. Models with no difference at all between update schemes, such as the Toll Pathway of Drosophila Signaling Pathway model [38], exhibit a kind of monostability in which only a single globally stable fixed point exists for each combination of source node values, regardless of update scheme; these models are highly ordered.

**FIG. 3.**
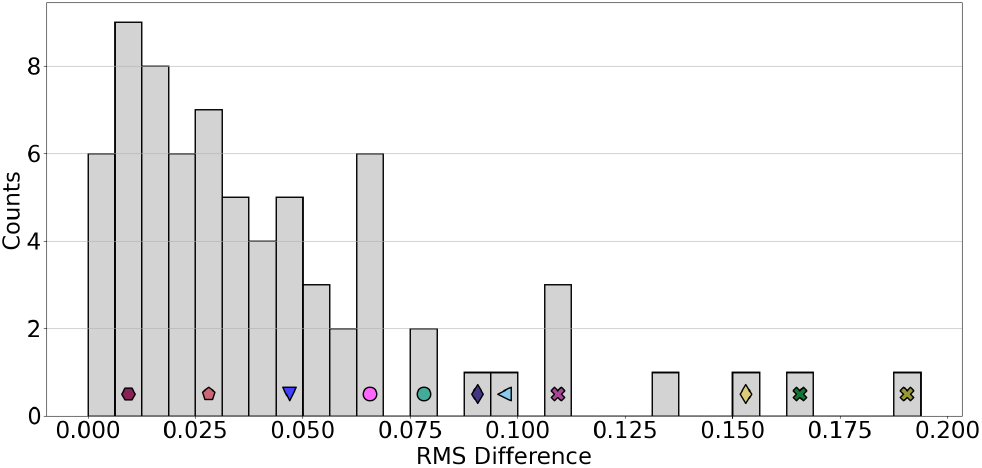
Distribution of update dependence in the Cell Collective. The root mean squared (RMS) difference between the node values when using synchronous or asynchronous update, as defined in Section II A, is shown. The peak near zero indicates a high degree of timing robustness in the Cell Collective models. Representative models are indicated by symbols according to Figure 1.

The Regulation of the L-arabinose operon in *Escherichia coli* 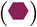 model [1] has two mechanisms through which population-level update scheme robustness arises. The first mechanism is trivial monostability, which applies in 11 of the 12 possible input combinations. In the last combination, as discussed in the previous section, the model has two point attractors in both update schemes and four additional cyclic attractors under synchronous update that arise from a positive feedback loop as in the example of Equation 1. As in that simple example, the symmetry of the positive feedback loop causes the average node values to be unaffected by the additional attractors.

Similarity between update schemes can also arise in more subtle ways; average node value similarity in the Metabolic Interactions in the Gut Microbiome 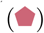 model [53] is driven by small, update-independent subnetworks. This model describes inferred interactions among 10 bacterial genera of the healthy gut microbiome, the pathogenic bacterium *Clostridium difficile*, and clindamycin antibiotic treatment. When clindamycin is present, the system reduces such that the attractor is determined by a complete subnetwork of three cooperative and self-sustaining bacterial genera. As a consequence, the basin of the attractor in which all three genera are present, representing more than 85% of the state space, is identical in the two updates (see panel B of FigureF.3 in Appendix F). The remaining state space is split between two very similar attractors in a manner that only weakly depends on update scheme. In the absence of clindamycin, only two nodes are free to vary and their average values depend mildly on update scheme.

In cases when timing robustness fails to emerge, the network typically has a large number of states that can evolve to more than one attractor in the asynchronous update. Under synchronous update, each of these states must deterministically evolve to only one attractor. When these states are heavily biased toward one attractor over another, the network can exhibit desynchronization sensitivity. An example of this is the Cortical Area Development 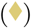 model [24], which aims to explain how interactions among a morphogen and four transcription factors lead to their characteristic expression pattern during mouse cerebral cortex development. The two poles of the cortex are represented by different initial conditions. The model uploaded to the Cell Collective was featured in [24] as a previously hypothesized model that does not recapitulate the expected biological result. We analyzed the model on the Cell Collective as well as one of the successful models reported in [24]. Both are bistable, with one attractor being much more likely than the other under synchronous update; under asynchronous update, the two attractors are more equally balanced (see Figure F.3 in Appendix F). The model in the Cell Collective is not successful under either update; the other model requires asynchronous update for success.

We caution that careful consideration of the underlying biology is always important when analyzing these models and selecting an update scheme, even when population-level average node values are fairly robust to timing perturbations. To illustrate this, consider the Apoptosis Network 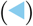 model [36] of Mai and Liu, which has an RMS difference in average node values that, though higher than the median, is quite low in absolute terms (near 0.1; see Figure 3). This model describes cancer cells’ decision between apoptosis and survival. Mai and Liu used synchronous update and report that both phenotypes are possible under each combination of growth factor and tumor necrosis source nodes. We confirm this and identify a three-node subnetwork that determines the phenotype.

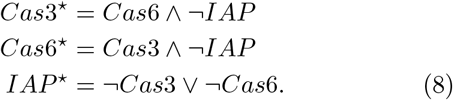

Apoptosis occurs when *Cas*3 = *Cas*6 = 1 and *IAP* = 0, while *Cas*3 = *Cas*6 = 0 and *IAP* = 1 lead to survival. Our analysis with cubewalkers found that the outcome of both the full model and this subnetwork strongly depends on update scheme: apoptosis is twice as likely under asynchronous update (see Figure F.2 in Appendix F). In the full model, changing the update scheme changes whether survival or apoptosis is more likely. Despite this dramatic difference, enough nodes in the network take the same value in both attractors that the network’s average node values overall are moderately robust to update scheme.

Though cases of update scheme dependence often highlight interesting regulatory mechanisms, we emphasize that population-level desynchronization robustness is by far more common in the Cell Collective. In combination with the results of the previous section, this points to an order in the average states of nodes that is hidden when these biomolecular networks are viewed as isolated entities but that is revealed when they are viewed as members of an ensemble.

### B. The effects of transient state perturbations

In the previous section, we discussed the effects of timing perturbations in Cell Collective models; we now consider the effects of transient node perturbations in which the state of a variable is temporarily altered. We emphasize the comparison between the short-term response measured by the Derrida coefficient (*δ*) and long-term responses measured by the quasicoherence (*q*), final Hamming distance (*h*^∞^), and fragility (*φ*), which are defined in subsection II D of the Methods and differ in how long-term changes to trajectories are quantified. We also consider the impacts of internal perturbations separately from those of environmental changes by considering two cases for all measures: perturbable and fixed source nodes, emulating a variable or static cellular context, respectively.

#### 1. The prevalence of source nodes in the models has a strong influence on trajectory separation

Previous studies did not consider the fact that the variables of Boolean network models fall into two qualitatively different categories: independent variables (represented by source nodes in the network) and variables whose values are determined by their interactions (represented by nodes with incident edges in the network). Source nodes are rare in most types of RBN ensembles. We determined (see Appendix E) that in any ensemble of finite random networks obeying widely used independence assumptions, on average more than 75% completely lack source nodes. This stands in stark contrast to the Cell Collective; only nine of the 72 models we studied are source-free, and the average number of source nodes in these networks is 4.94 (median 3, maximum 33) (see Figure E.1 in Appendix F for the full distribution). Note that these statistics and the distribution of the number of source nodes do not include constant nodes or source nodes for which only one value is ever considered in the analysis of a model’s original publication. The number of constant nodes in random models is much less tightly constrained than the number of source nodes, thus the frequency of constant nodes in our test ensemble could plausibly be obtained in random models (see Appendix E).

Dynamically and biologically, source nodes play an important role. In biology parlance, they often describe the cellular *context*, or configuration of the external environment and of intra-cellular mechanisms outside the scope of the model under study. Often, a change to the value of a source node represents an enormous shift in this context. This is because a change in the value of a source node is not a temporary dynamical perturbation, but a permanent alteration of the modeling context. Dynamically, this is reflected in the distribution of *δ* and *h*^∞^ in the Cell Collective ensemble (see Figure 4). When source nodes are perturbable in the synchronous update, we find that the distribution of *δ*_*s,p*_ peaks very close to 1. This corroborates previous observations[14, 55] in Boolean models of biological systems. However, an abundance of source nodes tends to increase *δ* in these models, in some cases dramatically, because the ultimate size of a perturbation that begins at a source node is always bounded below by one (in contrast, constant nodes tend to decrease *δ* because they are guaranteed to recover from any perturbation). Furthermore, many Cell Collective models are concerned with how signals, represented by source nodes, are processed by cells, meaning that–by design– such models tend to be sensitive to the values of these source nodes.

By isolating the effects of source nodes on the *δ*, we can begin to understand the degree to which the overall perturbation response in cellular systems is governed primarily by factors internal to specific functional modules (non-source nodes), or by the interplay between these modules and their environment (source nodes). When we restrict attention to the system’s response to internal perturbations only, we see that *δ* is no longer centered near one. Rather, the distribution shifts dramatically to the “ordered” regime (below 1). For example, the Metabolic Interactions of the Gut Microbiome 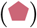 model has *δ ≈* 1 when source nodes are candidates for perturbation but only ≈ 0.39 when they are not. In the asynchronous case, defined in Equation 2, *δ* is more tightly clustered, but overall, *δ* shows very little dependence on the update scheme (see Figure F.4 in Appendix F for a direct comparison). This suggests that, on short time scales, the disorder that arises from node perturbations does not couple with the noise that arises from disruptions to update synchrony.

A few models do not follow the general trend and exhibit *δ* higher than 1. One example is the *Arabidopsis thaliana* Cell Cycle 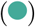 model [43] which has the highest value of *δ* (greater than 1.2 in both update schemes). This 14-node, source-free model has an abundance of regulators (average in-degree of 4.71), a significant percentage of which (42%) are negative regulators. The complexity of the regulation is likely the reason for the high observed initial separation of trajectories following an initial perturbation to a single node.

In the thermodynamic limit of random Boolean networks, there is a very strong relationship between *δ* and *h*^∞^. Whether or not this holds in the Cell Collective is investigated in Figure 4. The quadrants of the two panels of Figure 4 show whether the perturbation response indicates ordered or chaotic dynamics in the short-term or long-term. The short-term perturbation response of the models, as measured by *δ*, suggests ordered dynamics in the bottom two quadrants and chaotic dynamics in the top two quadrants. The long-term perturbation response, as measured by *h*^∞^, suggests ordered dynamics in the left two quadrants and chaotic dynamics in the right two quadrants. In the Cell Collective models, we observe a slight correspondence between *δ* and *h*^∞^ under synchronous update. No correspondence of *δ* and *h*^∞^ was found for asynchronous update (see Figure F.5 in Appendix F). It is somewhat expected that the correspondence between *δ* and *h*^∞^ would be stronger in synchronous update, where phase shifts within oscillatory attractors are always persistent. In contrast, phase shifts often decay in asynchronous update. When source nodes are not perturbable, *δ* serves as an upper bound for *h*^∞^in the ordered regime, and as a lower bound for *h*^∞^ in the chaotic regime (see Figure 4, right panel). For fixed source nodes, *h*^∞^ varies wildly when *δ* ≈1, reflecting an extreme sensitivity that is characteristic of systems near a phase boundary. Note that both *δ* and *h*^∞^ are skewed more toward the ordered regime than in the traditional approach of perturbable source nodes, shown in the left panel.

**FIG. 4.**
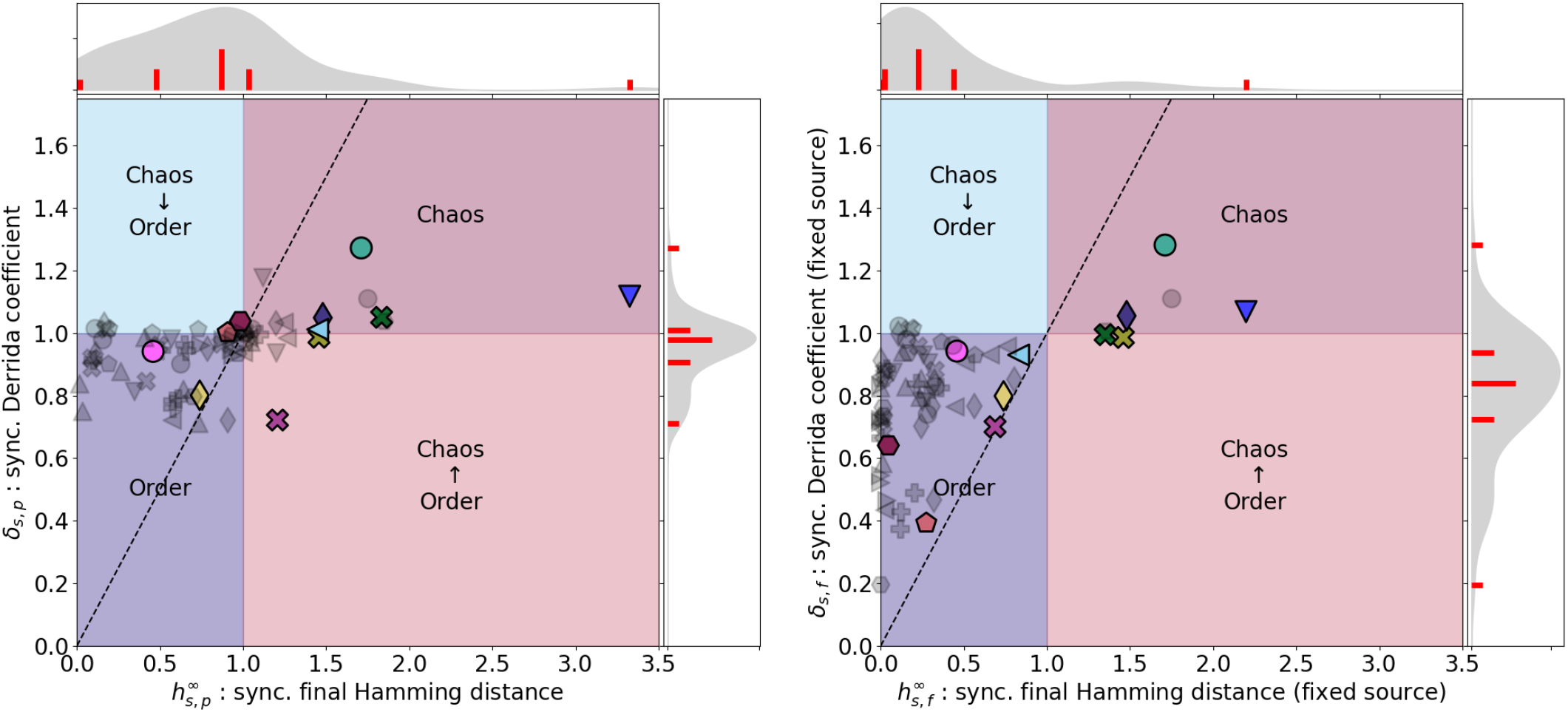
Short- and long-term perturbation responses in the Cell Collective measured in a phase-sensitive way. In the ordered regime (lower left quadrant) both short-term and long-term responses are below 1. In the chaotic regime (upper right quadrant) both short-term and long-term responses are above 1. The other two quadrants indicate cases of disagreement between the short-term and long-term responses. The short-term perturbation response *δ* has a slight correspondence with the long-term perturbation response under the specific setting when *h*^∞^ is monitored and synchronous update is used, in which the phase shifts are conserved. The relationship between short- and long-term responses is stronger when source nodes are fixed (right panel). The dashed line indicates the *y* = *x* diagonal. The symbols indicate the model categories and highlighted models as defined in Figure 1.

When source nodes are not perturbable, *h*^∞^ decreases dramatically for many models (see Figure F.6 in Appendix F). This is likely due to the large number of Cell Collective models that describe how functional modules integrate and respond to external signals, leading to a bias for source nodes with significant downstream effects. For example, as previously discussed, the Regulation of the L-arabinose operon in *Escheria coli* model 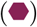 is monostable in most of its input configurations. This leads to very small *h*^∞^ when source nodes are not perturbable, despite the fact that this model has a slightly above-average *h*^∞^ when its source nodes are potential perturbation targets.

Models of functional modules with more complex internal dynamics, such as the Signal Transduction in Fibroblasts 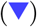 model [28] can also be greatly affected by source nodes. This model stands out in its high value of *h*^∞^, despite its only slightly elevated Derrida coefficient (*δ*_*s,p*_ = 1.12). This 130-node model describes the response of a specific cell type to 9 external signals (growth factors, cytokines, stress). These signals modulate the complex internal dynamics, but do not completely control them; thus the horizontal position of this model in 4 is further to the left when source nodes are fixed (right panel), but it remains the model with the highest *h*^∞^.

The Tumour Cell Migration and Invasion 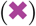 model [8] stands out in that it has a low value of *δ*, but a high value of *h*^∞^ in synchronous update when source nodes are perturbable. This model describes the processes necessary for cancer cell metastasis, including an epithelial to mesenchymal cell fate change, gain of motility, and the ability to invade the neighboring tissue. The model’s two inputs describe an internal signal (DNA damage) and an external signal from the cell’s microenvironment. The non-monotonic change in time of the Hamming distance persists in the input combination most relevant to cancer cells. One factor that contributes to a low *δ* (below 1) is the strong canalization of the model’s functions, which are biased heavily toward the “OFF” state. This causes many perturbed trajectories to immediately realign, resulting in a low *δ*. Though most trajectory pairs quickly align, those that do not tend to dramatically increase their separation, converging into very distinct attractors and resulting in a higher *h*^∞^.

#### 2. Perturbation response beyond trajectory separation

In this section, we use two measures introduced in Section II D, the quasicoherence *q* and fragility *φ*, to illustrate that it is difficult to alter the long-term dynamics of trajectories using small, internal perturbations. We demonstrate, in Figures 5 and 6, that careful comparison of the overall behaviors of perturbed and unpertrubed trajectories reveals a higher degree of order than is observable using traditional measures alone. The bulk of this section is devoted to uncovering the mechanisms that underlie this previously hidden order in specific models. We identify three key factors that give rise to disagreement between our new measures and traditional measures: i) the extreme potency of perturbations to source nodes, ii) the presence of oscillatory attractors that can result in phase-shifted trajectories with the same long-term behavior, and iii) higher sensitivity to update scheme in traditional measures.

The quasicoherence *q* describes the likelihood that a system undergoes a long-term phenotypic change in response to a small, transient perturbation. Higher *q* indicates a greater degree of phenotypic robustness (see II D). Note that the values of source nodes also contribute to the phenotype in this context, and so the effect of allowing source node perturbation is particularly pronounced for *q*. We find that overall, the distribution of *q* in the Cell Collective (Figure 5) is highly concentrated near 1 for the fixed-source case (see also Figure F.7 in Appendix F). This indicates that it is relatively difficult to alter the phenotype of a functional module within a cell by perturbing a single internal component. Indeed, no model has greater than a 60% chance to change quasiattractor due to perturbation to a random node; when source nodes are excluded from the set of perturbable nodes, this bound drops to just over 20%. An example of low quasicoherence is the Cortical Area Development Network 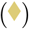 model [24], which has two attractors; the symbol lies on the diagonal because this model has no source nodes.

**FIG. 5.**
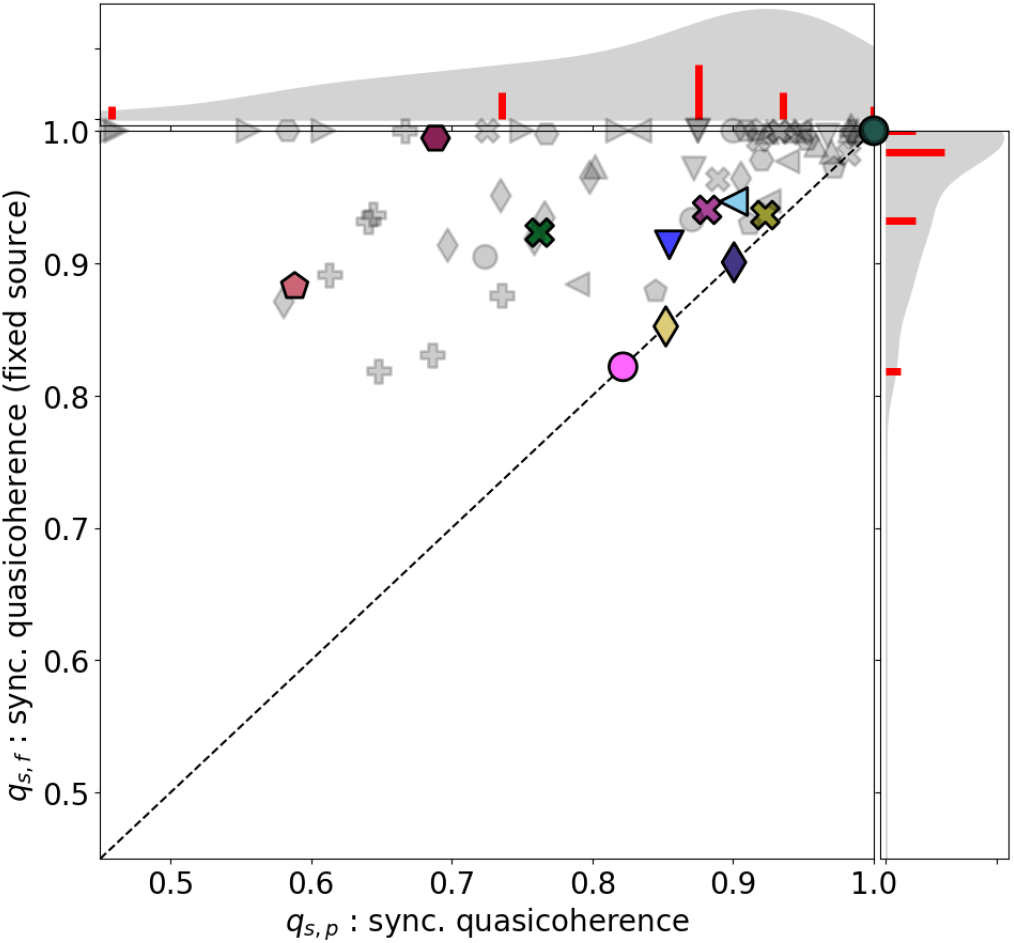
Scatterplot of the synchronous quasicoherences of the Cell collective models when source nodes are (x axis) or are not (y axis) candidates for perturbation (the asynchronous distribution is available in Figure F.7 of Appendix F). When the values of source nodes are fixed, the quasicoherence values are tightly clustered around 1, indicating a high degree of phenotypic robustness. The symbols indicate the model categories and highlighted models as defined in Figure 1.

The distribution of *q* in the Cell Collective is fairly robust to update scheme, though there are exceptions. For example, note that Cell Cycle Transcription by Coupled CDK and Network Oscillators 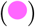 model has relatively low quasicoherence in the synchronous update, but a maximal quasicoherence in the asynchronous update (see Figure in Appendix F). The difference arises because the asynchronous update gives rise to only a single attractor (a steady state) while the synchronous update gives rise to an additional oscillatory attractor. In this case, the timing perturbations have interfered with the node perturbations in the system by destroying an attractor that is required for long-term separation of trajectories. The fragilities *φ* of the Cell Collective models also exhibit a distribution that is generally robust to the update scheme, and a shift to lower values when source nodes are not candidates for perturbation (see Figure F.8 in Appendix F).

Separate from quantifying whether or not a perturbation induces a change in phase-shift-corrected long-term behavior (via *q*), we also quantify the magnitude of such changes using *φ*. Figure 6 summarizes the relationship between *δ* and *φ* under synchronous update with fixed source nodes. Note that only two models exhibit chaotic long-term perturbation response once source nodes and phase shifts are accounted for, and the vast majority of the models are firmly in the ordered regime of the *φ* distribution. In contrast, the traditional measures place the majority of the models close to the critical boundary between the ordered and chaotic regimes, and also place several models in the chaotic regime (left panel of Figure 4). We found no correspondence of *δ* with *φ* regardless of the manner of update or the perturbability of source nodes. Furthermore, unlike in the case of *h*^∞^, the *φ* distribution shows little dependence on the choice of update scheme. (See Figure F.5 in Appendix F for a comprehensive figure combining Figures 4 and 6 with five other similar plots). This suggests that the ability of *δ* to predict long-term perturbation response is sensitive to phase-shifts and can overestimate the disruption a perturbation is likely to cause to a system’s phenotype.

**FIG. 6.**
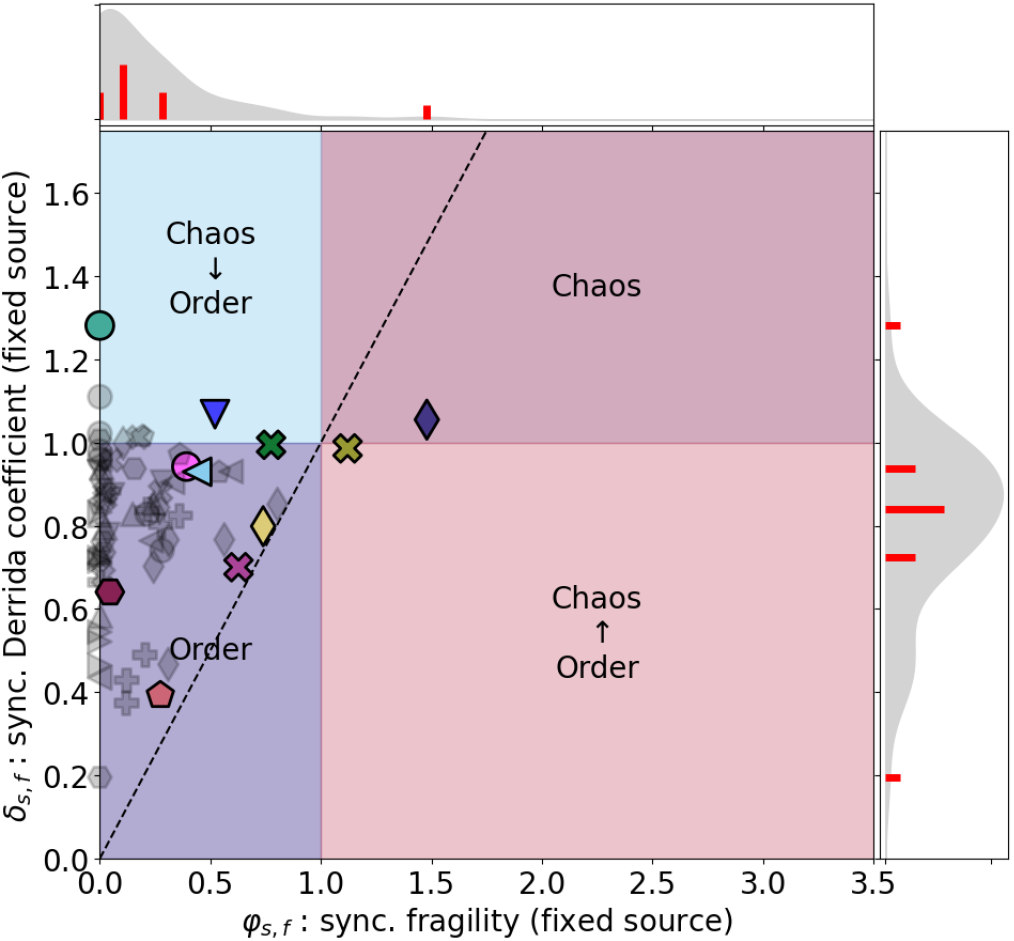
Short- and long-term perturbation responses in the Cell Collective measured in a phase-insensitive way. In the ordered regime (lower left quadrant) both short-term and long-term responses are below 1. In the chaotic regime (upper right quadrant) both short-term and long-term responses are above 1. The other two quadrants indicate cases of disagreement between the short-term and long-term responses. In contrast with the traditional approach depicted in the left panel of Figure 4, this figure illustrates perturbation response when source nodes and phase shifts are accounted for. Most models show an ordered perturbation response when these factors are taken into consideration. The symbols indicate the model categories and highlighted models as defined in Figure 1.

As we illustrate with several examples below, it is often possible to reveal a robust order in apparently chaotic perturbation responses of specific functional modules by carefully analyzing the patterns of oscillation that perturbed trajectories undergo.

As highlighted previously in Figure 4, the Signal Transduction in Fibroblasts 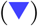 model [28] has a very high value of *h*^∞^ in the synchronous update (>3 when the source nodes can be perturbed and 2.3 when they cannot), and *δ* only slightly above one. Asynchronous update decreases *h*^∞^ but preserves the “chaotic” classification (see Figure F.5 in Appendix F). We found that this model has an exceptionally rich repertoire of quasiattractors, with an average of more than 700 quasiattractors for each of 512 source node combinations. The vast majority of these quasiattractors correspond to oscillations. Given this richness of phenotypes, large responses to perturbations may be expected. Despite this, *φ* is less than one in both update schemes in this model when source nodes are fixed, meaning that at the phenotype level, perturbations to individual nodes eventually decay on average. In other words, the majority of the perturbation response observed through the lens of *h*^∞^ is due to the effect of shifting the phase of a trajectory without altering its phenotype. The Aurora Kinase A in Neuroblastoma 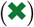 model is a smaller model that exhibits similar behavior.

The *Arabidopsis thaliana* Cell Cycle 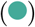 model [43] is also in the chaotic regime when synchronous update is used to compute *δ* and *h*^∞^ (Figure 4), but a closer look reveals a robust phenotype. The original article reported an 11-state cyclic attractor under synchronous update, which recapitulates the phases of the cell cycle, and in which all the 14 nodes oscillate. This model’s response to an initial perturbation to a single node is the highest observed (*δ >* 1.2 in both update schemes). In the synchronous update, this initial separation persists, and even grows somewhat in the long term (reaching an average of over 1.7). Because there is only one attractor in this system, and because synchronous attractors are always simple cycles, this separation is due to a phase shift; indeed, the fact that the synchronous fragility of this model is zero reinforces this (Figure 6). In the asynchronous update, both the fragility and the final Hamming distance are zero, indicating that this model exhibits a long-term order under the asynchronous update that is not detected by *δ*. The difference in long-term separation in the two updates reflects the fact that phase shifts are always permanent in the deterministic synchronous update, but can be temporary in the asynchronous update if there is an order of update that causes two trajectories in the same complex attractor to intersect. Indeed, there is a general tendency for a smaller final Hamming distance under asynchronous update than under synchronous update (see Figure F.6 in Appendix F). Furthermore, Figure F.9 in Appendix F suggests that phase-shifting behavior of the *Arabidopsis thaliana* Cell Cycle 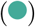 model is a common phenomenon; the final Hamming distance is always larger than or equal to the fragility in both update schemes, with an especially prominent difference in synchronous update.

There are two models that stay in the chaotic regime according to both *h*^∞^ and *φ*, the Human Gonadal Sex Determination 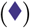 model [46], and the Colitis-associated Colon Cancer 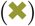 model [35]. These two are the only models with *φ >* 1 when source nodes are not candidates for perturbation. The Human Gonadal Sex Determination model 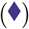 describes the gene regulatory network that controls the differentiation of the gonadal primordium towards testes or ovaries in the early stages of embryonic development. The original article reported three point attractors; in addition to the two expected ones, each with a basin of almost 50% under synchronous update, there is a third attractor, corresponding to disgenetic testes, whose basin is less than 1%. We find that under asynchronous update the basin of the two expected attractors decreases and the basin of the third attractor increases. The Colitis Associated Colon Cancer model 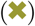 integrates the signaling pathways that underlie inflammation-associated tumorigenesis. This model has four attractors, three of which have a sizable basin, but no basin dominates the state space. The existence of three or more attractors is a major factor in the high fragility of these models. Fragile models such as this are characterized by basins of attraction with attractors that differ in many nodes. When a node of the system is perturbed, the system has a tendency to enter a different basin of attraction, causing its converged average node values to be substantially different than those of the unperturbed trajectory.

A four-node reduced version of the Human Gonadal Sex Determination model 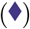 illustrates this property.

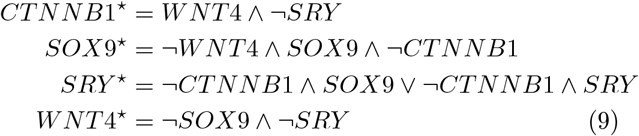

This reduced model has three attractors, one of which has a basin of attraction much larger than the others (11 states vs 2 and 3 states). The two attractors with smaller basins of attraction are highly fragile; a perturbation to a single node has a 75% chance of altering the attractor basin in four out of five of these states, and a 50% chance of doing so in the fifth state. Though these attractors have small basins, collectively they make up just under a third of the state space. See panel C of Figure F.2 in Appendix F for a detailed visualization of the fragility of this reduced system.

In summary, our analysis of the Cell Collective models using our newly introduced measures of quasicoherence and fragility reveals that most of them are phenotypically ordered for both update schemes considered. With these measures, we uncover nontrivial perturbation recovery on long timescales even in putatively chaotic perturbation responses captured by the final Hamming distance, and we identify key mechanisms behind phenotypic fragility and robustness.

## IV. DISCUSSION

One of the conjectured hallmarks of complex biological systems is that they sit somewhere between rigid order and hypersensitive disorder. For example, a yeast cell must be able to adjust its metabolic phenotype in response to external cues such as oxygen availability, and to internal cues that operate downstream of cellular mechanisms involved in processing environmental signals. At the same time, the yeast cell must not chaotically switch between metabolic pathways in response to small fluctuations in external conditions or in response to noise in its internal regulatory processes. From an evolutionary perspective, some degree of phenotypic mutability confers adaptability to a population; too much leads to lack of evolvability or even population collapse [10]. It has been argued that in living systems, there is often a sharp boundary between these regimes, and the cusp of this boundary is the ideal place to balance these competing needs [6, 16, 17, 44, 48, 51]. Indeed, in simple random models that resemble biomolecular regulatory systems, this appears to be the case [4, 16, 17, 61]. The argument is further bolstered by the fact that real-world models of specific within-cell functional modules share some properties exhibited by these simple random models in the critical regime [12, 27, 30, 37].

But these real-world models are not random; for instance, they exhibit a higher degree of canalization and functional redundancy [12, 22, 39], as well as a higher occurrence of source nodes (as demonstrated here). Of course, it is well-known that these models are non-random, and researchers are typically careful to acknowledge the caveats this entails. For example, Kauffman considers the question of random network assembly in some depth from a biological perspective [48]; Moreira and Amaral give a rigorous treatment of the implications of nonergodicity and canalization in Boolean ensembles [39]; Zañudo and colleagues give a careful treatment of the underlying assumptions of randomness and their implications [61]; and we ourselves have discussed the potential pitfalls of applying techniques designed for random networks to non-random networks in previous work [12, 37]. The Derrida coefficient [16, 17], or its close cousin, the network sensitivity [51] are superb tools in the setting in which they were developed: synchronously updated random models. In that setting, they offer a computationally simple way to determine the short-term and long-term response of the system to perturbations. Even in non-random models, these tools remain valid for exploring the short-term perturbation response, but more sophisticated measures are required for studying their long-term dynamics in response to perturbations.

The traditional approach to directly quantifying the long-term response to perturbations is to measure what we have called the final Hamming distance. This measure provides valuable information about the asymptotic separation of perturbed and unperturbed trajectories, but fails to account for time-shifts. By considering whether perturbed and unperturbed trajectories differ in ways that are in principle observable under typical experimental settings, the new measures we introduce provide a phenotypically grounded way to quantify the ultimate impact of a perturbation. Our analysis shows that the responses to internal perturbations that have been previously associated with criticality are usually either more transitory than initial perturbation growth may suggest or become phenotypically irrelevant in the long term. In fact, in the studied experimentally-supported, non-random models we uncover much greater robustness to perturbation, especially in their long-term effects, than the criticality hypothesis implies.

Though such orderly behavior of functional modules has been overlooked, indeed hidden by the typical measures of criticality used, it is not altogether surprising. For example, it is fundamental to Kauffman’s thesis that orderly behavior can arise naturally from RBNs [32, 48] and may play a key role in the evolution of epigenesis. More recent work [52] has analyzed microarray time-series data to suggest that eukaryotic cells do not lie in the chaotic dynamical regime. Particularly at the scale of individual functional modules, we would expect a high degree of reliability in task execution under most perturbations. For example, to effectively balance photosynthesis efficiency with water conservation, the regulatory mechanism of stomatal guard cells in plant leaves must reliably respond to stress hormones produced by other modules in the plant’s regulatory network. Indeed, we observe that in the Guard Cell Abscisic Acid Signaling model [34] and the Stomatal Opening Model [20], the fixed-source fragility is quite low (see Appendix G). In contrast, the traditionally-used Derrida coefficient suggests functional modules near or in the chaotic regime. We interpret this to suggest that small errors in signal transduction may lead to large initial deviations in these systems, but that eventually, these errors are corrected in most cases. In the context of cell differentiation, Waddington [58] argues for a kind of long-term developmental robustness referred to as canalization; once committed to a cell fate, it is expected that a stem cell is not easily diverted from its specialization. We observe this in various development and differentiation models, such as the Lymphoid and myeloid cell specification and transdifferentiation model [9]. In this model, the short-term perturbation response suggests criticality (*δ*_*s,p*_ = 1.02), but a long-term view reveals that initially divergent perturbed trajectories are canalized toward the fate of their unperturbed counterparts in most cases (*q* = 0.9_*s,p*_, *φ*_*s,p*_ = 0.16).

The new measures we introduced to characterize this robustness or phenotypic order allow us to distinguish process delay from phenotype differentiation (*h*^∞^ vs *φ*), and to separate smoothly varying distance in -omics space from “all-or-nothing” phenotype differences (*φ* vs *q*). These measures are computationally expensive to estimate, and until now, their estimation on ensembles of large models (more than a few dozen nodes) has been prohibitive. Here, we have addressed this challenge by developing cubewalkers, a highly-parallel GPU-based simulation toolkit. Our analysis showcases its capacity for comprehensive calculation of long-term perturbation dynamics in real-world Boolean networks with hundreds of nodes or more. Together with traditional measures, our new approaches offer a more holistic way to study the dynamical response of living systems to noise and perturbation.

Though our analysis suggests that the criticality of experimentally supported Boolean models of biomolecular functional modules has been overstated, we emphasize that this work is not the nail in the coffin of the “edge of chaos” hypothesis. Rather, it suggests that living systems do not exhibit critical behavior *at the scale of functional modules*. This leaves ample room for critical behavior to emerge at larger scales via the coupling of various functional modules. Indeed, previous work by Balleza and colleagues ([6]) suggests cell-scale critical perturbation response in two full-genome regulatory networks with experimentally constrained topology and random regulatory functions, though the authors do not consider phase shifts in their analysis. We conjecture that individual subsystems of a cell are highly ordered, but they connect in networks that may give rise to more adaptive behavior. The large differences in perturbation response we have observed depending on the treatment of source nodes (which are exceedingly rare in traditional RBN models) supports this conjecture because it allows for larger perturbation responses in networks of highly ordered functional modules coupled at their source nodes. In critical RBNs, one may view the nodes themselves as ordered subsystems. In real biological systems of many variables, a multi-scale, modular structure is expected [50]. Thus, it is possible that order persists up to larger scales in biology than it does in random models. More thorough examination of criticality and perturbation response across regulatory scales is needed to test our conjecture, which motivates the future development of sufficiently data-constrained multi-scale models.

Despite our finding that the Derrida coefficient is not a good predictor of phenotypic robustness, we do not suggest that it is without merit in models of specific functional modules. Instead, we merely caution that it must be carefully interpreted as an indicator of *immediate* response to perturbation only and should be studied in conjunction with long-term response measures, such as those we have developed here. We do however suggest that careful consideration be made to the biological interpretation of source node perturbation in the context of the particular network being considered. Generally, we advise that perturbation of these nodes be handled separately from perturbations to other nodes in the network.

We have also studied timing perturbations in these systems by considering the effect of update scheme on various dynamical properties. Many update schemes exist for Boolean networks, such as the most permissive Boolean network framework of [45], random order update [3], or various update schemes that make use of a continuous time parameter such as is used in MaBoSS [54]. We focused on the synchronous update and the asynchronous update, which are the most frequently used and are the two opposite extremes of the spectrum from deterministic timing coherence to completely stochastic event timing. It is well established that the update scheme can have a dramatic impact on the attractor dynamics of Boolean networks (see e.g., [47]). In the models considered here, the average behavior of individual system components is typically quite robust to update scheme, but in a few models there is a dramatic difference in the biological interpretation of the individual trajectories that are possible in one update scheme or the other. In the examples we have examined here where this is the case, there are attractors that exist in the synchronous update but which are absent in the asynchronous update. In all such cases, the attractors were motif-avoidant, i.e. they did not fall into any minimal trap space [47] (sometimes these are called unfaithful attractors [33]). In these examples, delay nodes played a prominent role in the behavior of the model under synchronous update. We generally found that models appear more ordered in the asynchronous update, for example, via the destruction of synchronous attractors. Most dramatically, the median value of *h*^∞^ for fixed source nodes is approximately 43% higher in the synchronous update than in the asynchronous case. We conjecture that noise in the update timing can suppress the phase-dependent effects of node perturbation. Indeed, while two phase-shifted oscillating trajectories can never realign in the synchronous update, eventual realignment is likely under the asynchronous update. Thus, the long-term response to node perturbations becomes biased toward extinction in the asynchronous update as measured by *h*^∞^ (see Figure F.6 in Appendix F). In contrast, because *q* and *φ* inherently account for phase-shifts in perturbed trajectories, they are much less sensitive to update scheme (see Figures F.7 and F.8 in Appendix F).

We have illustrated the overall patterns observed in the experimentally supported model ensemble by carefully examining the dynamics of specific examples and considering dynamical behavior in the context of their intended biological modeling goals. This has highlighted that the rich diversity of biological function is not easily distilled to a few statistical properties. Some functional modules have dynamics that almost trivially follow from the configuration of their inputs, while others modules are highly multi-stable with long-term dynamics that depend strongly on initial conditions and internal timings. In the search for unifying principles in biology, it is important to acknowledge that biology is messy and that functional context matters—especially in the study of specific subsystem models. In other words, living systems are complex, open systems. While there are important general conclusions we can draw, the differences between biomolecular systems can be just as interesting as their common properties. In that spirit, we show that functional modules in biomolecular systems typically exhibit robust phenotypes,while highlighting the diverse mechanisms through which this hidden order can arise. The observed order, as a phenomenon of experimentally supported models, has been hitherto obscured by the lack of dynamical measures that can quantify it and the computational challenges of measuring the dynamics with sufficient detail, an obstacle we overcame in the present work.

We hope that as computational biology continues its second half-century, unprecedented computational power allows deeper exploration of the interplay between order and chaos in living systems, and helps uncover the unique biological circumstances that enable it.

## ACKNOWLEDGMENTS

We thank Dr Jorge Gómez Tejeda Zañudo for his helpful advice regarding the framing of our results. This work was funded by NIH National Library of Medicine Program grant 01LM011945-01 to L. M. R., the Fundação para a Ciência e a Tecnologia grant 2022.09122.PTDC to L. M. R., and NSF grant MCB1715826 to R.A.

## DATA AVAILABILITY

The cubewalkers library is open source and available at https://github.com/jcrozum/cubewalkers. Data analysis scripts and raw data are available at https://github.com/troonmel/cubewalkers-analysis. All other materials are provided in the appendices.

## Appendix A: Benchmarks

In this section, we present benchmarks comparing the cubewalkers software to two competing software packages: cana and booleannet.

**TABLE A.1.** Average (amortized) run time per simulation time step for each method. The fastest method (cubewalkers) for the two hardware configurations are bolded. Note that cube-walkers on consumer hardware outperforms parallel adaptations of other tools running on specialty high-performance computing hardware.

| software | hardware | time (μs) |
| --- | --- | --- |
| CUBEWALKERS | consumer | **0.11** |
| CUBEWALKERS | specialty | **0.067** |
| CANA (serial) | consumer | 40 |
| CANA (parallel) | specialty | 1.1 |
| BOOLEANNET (serial) | consumer | 1300 |
| BOOLEANNET (parallel) | specialty | 37 |

## Appendix B: Convergence of average node values

The number of walkers were selected to ensure a standard deviation of less than 0.01 for each dynamical measure computed. The minimum simulation count of *W* = 2, 500 was used in the calculation of average node values. The convergence of these values as a function of *W* is shown in Figure B.1. For Derrida coefficient calculation, a value of approximately *W* = 100, 000 was used (*W* = ⌊100 000*/N⌋* × *N*); for measuring long-term perturbation spread, a value of *W* = 2, 500 was used for each node targeted for perturbation (for a total of 2, 500 ×*N* simulations each, resulting in *W* = 800, 000 in the largest model considered).

**FIG. A.1.**
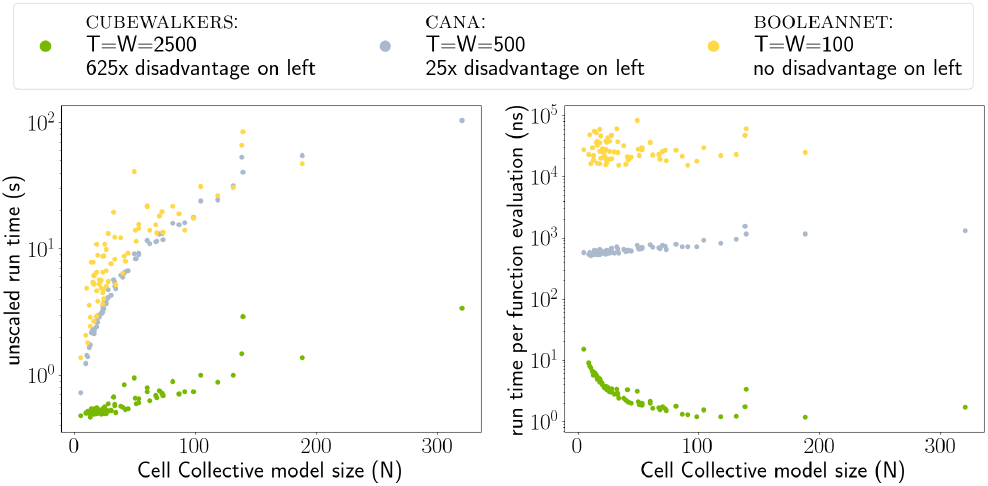
Performance comparison of CUBEWALKERS,CANA, and BOOLEANET on consumer hardware. 72 Cell Collective models were run using each tool using synchronous update. Timings were generated on a PC with an AMD Ryzen 5 3600X CPU at 3.8GHz and a 2560 CUDA-core 1605MHz NVIDIA 2070S GPU. Default methods were run without additional parallelization. For the CUBEWALKERS tests, 2500 time steps and 2500 walkers (initial conditions) were used; for CANA, 500 time steps and 500 walkers were used; and for BOOLEANET, 100 time steps and 100 initial conditions were used. Thus, for each network, CANA computed 5x as many time steps for 5x as many initial conditions as BOOLEANET for an overall disadvantage of 25x. Similarly, CUBEWALKERS computed 5x as many time steps for 5x as many initial conditions as CANA, for a 25x disadvantage relative to CANA and a 625x disadvantage relative to BOOLEANET. The raw time to complete these tasks is plotted in the left panel, where we observe that cube-walkers consistently finishes its tasks an order of magnitude faster than the other methods, despite the fact that it has been given significantly more computational work. In the right panel, the average computation time per network node per time step per initial condition in these trials is plotted; this corresponds to the average (amortized) time to evaluate and apply an update function to a node. Here, we see that these amortized evaluations occur on the order of nanoseconds for CUBEWALKERS, while they occur on the order of microseconds for CANA and hundreds of microseconds for BOOLEANET.

The question of how many time steps are required to have a reasonable expectation of average node value convergence is more complicated. There are two reasons for this: i) convergence time is highly model-dependent and as the systems considered are generally not ergodic, the average node values may converge into oscillatory behavior. Thus, there are two parameters that need to be considered: a “burn-in” time *T*_*b*_, and an averaging time window size *T*_*w*_, for a total simulation time of *T* = *T*_*b*_ + *T*_*w*_. We fixed *T*_*b*_ = 50*N* + 1 000, so that at least 1 000 updates are performed and each node is updated more than 50 times on average in the asynchronous update during the burn-in stage. We then varied *T*_*w*_ and evaluated the convergence of the average node values by comparing the values calculated in four sub-windows: *T*_*wi*_ = [*T*_*b*_ + *iT*_*w*_*/*5, *T*_*b*_ + (*i* + 2)*T*_*w*_*/*5], for *i* = 0, 1, 2, 3. For each network in the Cell Collective, we computed the absolute difference in average node values for each of the six pairs of these four sub-windows, and identified the largest absolute difference across all six comparisons for each node. Convergence quality is assessed by computing the largest of these values across all nodes. Based on this analysis, we chose to use a value of *T*_*w*_ = 5*N* + 5 000 for most models. Three models took an unusually long number of time steps to converge due to the complexity of their attractors; for these we set the number of time steps manually: (*T*_*b*_, *T*_*w*_) = (5 000, 25 000) for “Arabidopsis thaliana Cell Cycle” (*N* = 14), (*T*_*b*_, *T*_*w*_) = (5 000, 25 000) for “Guard Cell Abscisic Acid Signaling” (*N* = 44), and (*T*_*b*_, *T*_*w*_) = (50 000, 100 000) for “Signal Transduction in Fibroblasts” (*N* = 139). The largest absolute difference of average node values between any two time sub-windows

*T*_*wi*_ and *T*_*wj*_ across all nodes in all networks was approximately 0.0004 in the synchronous update and 0.0066 in the asynchronous update. Summing the largest difference for each node gives a maximum of 0.0039 and 0.0881 for synchronous and asynchronous update, respectively, across all networks in the Cell Collective. The actual computed quantities aggregate many nodes and average over a time window 2.5 times larger than any *T*_*wi*_; thus, in practice, they have errors much lower than this very conservative upper bound. We are therefore confident that simulating each network for *T* = 55*N* + 6 000 time steps and averaging node values over the last *T*_*w*_ = 5*N* + 5 000 time steps is sufficient for computing the average behaviors of nodes in almost all models in the Cell Collective.

**FIG. A.2.**
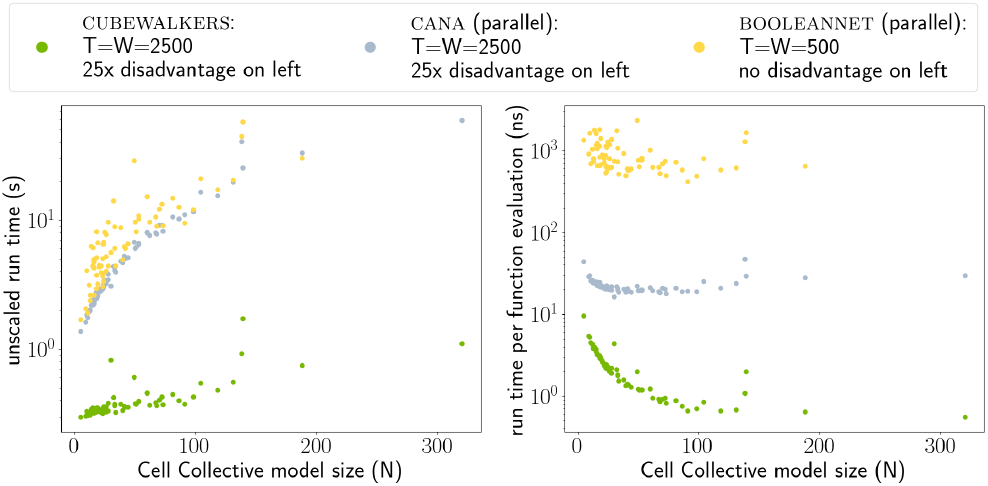
Performance comparison of CUBEWALKERS, CANA, and BOOLEANET on a high performance computer. Cell Collective models were run using each tool using synchronous update. Timings were generated using a workstation with two AMD EPYC 7542 CPUs (32 cores and 64 threads each) at 2.9GHz and two 10,752 CUDA-core NVIDIA A6000 GPUs with 48GB of GDDR6 memory (only one GPU was used for the benchmarks). For the CUBEWALKERS and CANA tests, 2500 time steps and 2500 walkers (initial conditions) were used; for BOOLEANET, 100 time steps and 100 initial conditions were used. For CANA and BOOLEANET, initial conditions were simulated in 128 parallel threads. On specialized hardware taking full advantage of parallelism, we see that the performance gap between CUBEWALKERS and the other methods is narrowed compared to the performance gap on consumer hardware. Nevertheless, the gap remains considerable.

**FIG. B.1.**
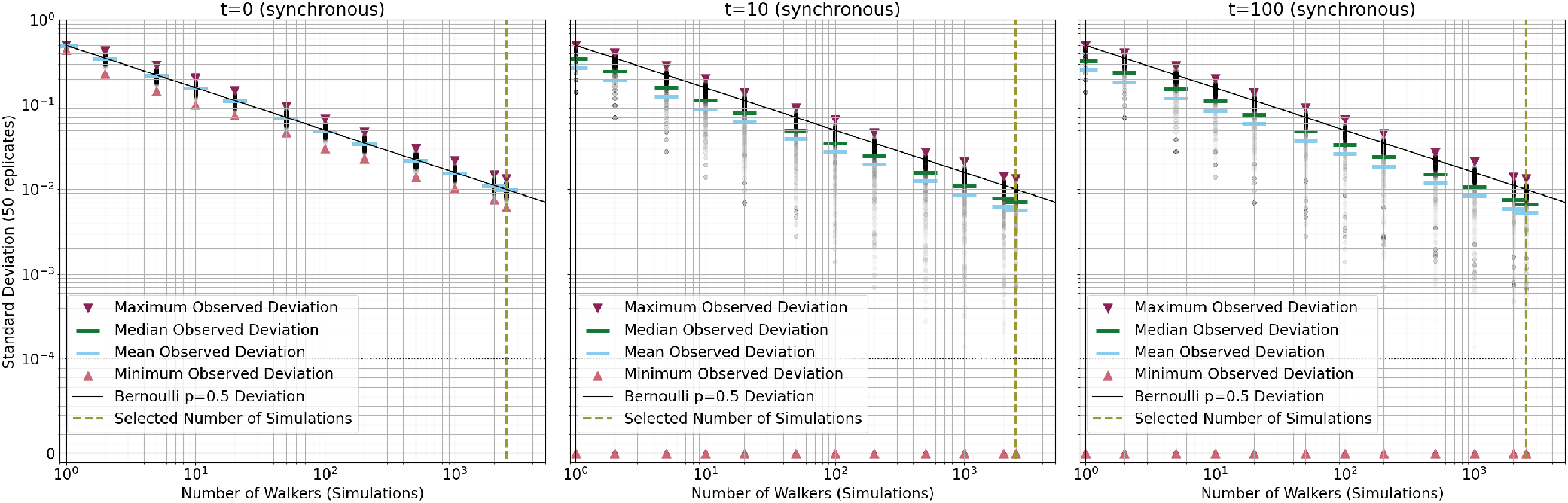
Standard deviation in the node average values as the number of walkers increases for the Cell collective models. The three panels correspond to three different stages of the model simulation. The observed standard deviations agree well with the expectation based on Bernoulli random variables (continuous line). We chose the number of walkers such that the standard deviation is less than 0.01 (dashed vertical line).

## Appendix C: The relationship between two dynamical measures: fuzzy quasicoherence and fragility

The quasicoherence measure treats trajectories that converge to the same quasiattractor as equivalent, even if they converge to different attractors within that quasiattractor. We introduce the fuzzy quasicoherence, a modification of the quasicoherence such that it becomes sensitive to the similarity of attractors but retains phase-insensitivity.

This is achieved by replacing the *Q* function with a “fuzzy” version that considers the absolute difference between *X*(*t*) and *X*^(*¬i*)^(*t*)). This gives rise to the fuzzy quasicoherence, 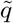:

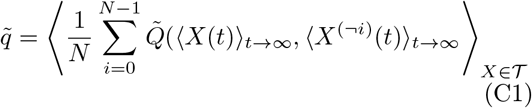

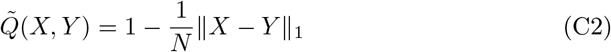

Note the similarity with both *q* and *h*^∞^. Compared with *q*, the formula for 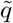 replaces the *Q* function with 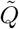, which, like *Q*, is 1 if the inputs are equal and 0 if the inputs are maximally different in each entry, but which can interpolate between 0 and 1. The ability to interpolate between the extremes of *Q* allows 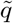 to account for whether quasiattractors are similar or different, and it also allows 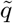 to account for attractors within the same quasiattractor that have different average node level behaviors. Compared with *h*^∞^, 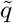 can be viewed as a rescaling with a slightly modified averaging scheme.

The fragility is related to the fuzzy quasi-coherence by the relationship 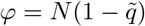.

## Appendix D: Detailed discussion of the Aurora Kinase A in Neuroblastoma model

The Aurora Kinase A in Neuroblastoma 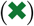 model developed by Dahlhaus et al. [13] explores the role of the Aurora Kinase A protein in the cell cycle of neuroblastoma cancer cells. Dahlhaus et al. used synchronous update and reported three families of attractors: a point attractor corresponding to the G0 checkpoint, a threestate cycle describing cells proceeding faithfully through mitosis, and a three-state cycle corresponding to cells with defective mitosis, respectively. Aurora Kinase A is off in the G0 point attractor, expressed and active in the faithful mitosis attractor, and oscillates in the defective mitosis attractor. Defective mitosis leads to mitotic catastrophe and cell death via mechanisms outside the model, and is desirable in the context of neuroblastoma. Dahlhaus et al. find that constitutive activation of Greatwall/MASTL stabilizes Aurora Kinase A, avoiding mitotic catastrophe and leading to poor prognosis; this was clinically confirmed by analyzing gene expression profiles of neuroblastoma patients. We found that the defective mitosis attractor requires synchronous update. In asynchronous update, only attractors corresponding to the G0 checkpoint and faithful mitosis exist. This also leads to population-level differences in this model: Aurora Kinase A is active significantly more often under synchronous update than under asynchronous update, yielding a higher average expression level of Aurora Kinase A in a cell population. This model can be reduced to the system

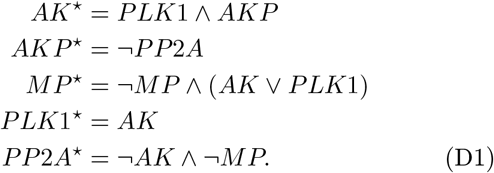

Here, AKP and AK represent the presence and activity of the Aurora kinase A, respectively; PP2A and PLK1 are important cell cycle proteins, and MP represents the physical processes of mitosis. As in the full model, this reduced system has synchronous-update attractors corresponding to the G0 checkpoint and faithful and defective mitosis; the last of these vanishes in asynchronous update, leading to differences in the average activity of Aurora kinase A. Notably, the synchronous behavior is sensitive to the existence of the intermediary node AKP: if AKP and AK are merged, the synchronous update yields similar results to the asynchronous update, which is insensitive to this merger. This shows that the defective mitosis attractor is dependent on a delay between PP2A activation and its effect on AK. Because delays are intrinsically stochastic in the asynchronous update, this delay dependency explains why defective mitosis cannot be sustained under asynchronous update.

## Appendix E: Source nodes and constant nodes are rare in RBNs

Source nodes are rare in most types of RBN ensembles. To illustrate this, consider an RBN ensemble with a specified in-degree distribution, *P* (*k*), and assume that a node with a given in-degree has its regulators chosen uniformly at random. We also assume that regulatory functions are chosen as in the N-K model with bias *p*. In such a random model, the probability that a node with in-degree *k* self-regulates is *k/N*, for a network of *N* nodes. The probability that the source update function (e.g., *f*_*i*_(*x*) = *x*_*i*_) is chosen is 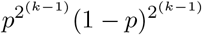 because for the half of the 2^*k*^ possible inputs in which *x*_*i*_ = 1, an output of 1 must be chosen, while for the other half, 0 must be chosen. Therefore, the probability that a node with *k* regulators and bias *p* is a source node is

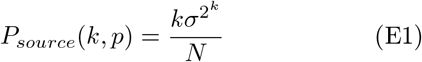

where we have used the bias variance, *σ*^2^ = *p*(1−*p*), to simplify the expression.

Thus, the probability that a specific node is a source node is 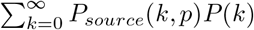. By assuming that node properties are generated independently, the expected number of source nodes can be calculated by multiplying by *N* :

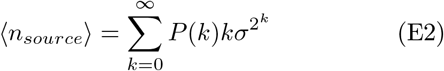

Notably, this expression is independent of *N*. This is because there are two competing effects as the network size grows that exactly cancel out on average: i) with more nodes, there are more potential source nodes, and ii) with more nodes, there are more potential regulators for each node, make it less likely that a node selects itself as a regulator.

We now put an upper bound on ⟨ *n*_*source*_⟩. The largest *σ* can be is 1*/*2, which is obtained for *p* = 1*/*2. This allows us to write 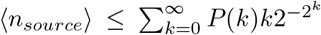.The expression 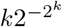 is maximized for *k* = 1. Substituting this provides a numerical upper bound on the expected number of source nodes

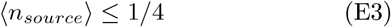

Because the expected number of source nodes is bounded above by 1*/*4, and because the number of source nodes in any finite network must be a nonnegative integer, we expect that in any ensemble of finite random networks (generated according to the assumptions above), more than 75% completely lack source nodes. This stands in stark contrast to the Cell Collective; only nine of these 72 models are source-free, and the average number of source nodes in these networks is 4.94 (median 3, maximum 33). (see Figure E.1)

A similar calculation can be performed to determine the expected number of constant nodes in these models. The probability that a node with *k* regulators has an update function equal to 1 is 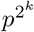 (because an output must be chosen for all 2^*k*^ input configurations). Similarly, the probability that this node has the update function 0 is 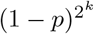. Thus, the expected number of constant nodes is

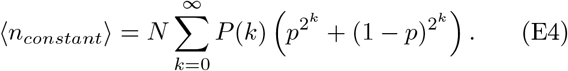

For *p* = 1 or *p* = 0, all nodes are constant; for *p* = 0.5, the fraction of constant nodes is minimized and can be made arbitrarily small by weighting the in-degree distribution toward higher *k*.

**FIG. E.1.**
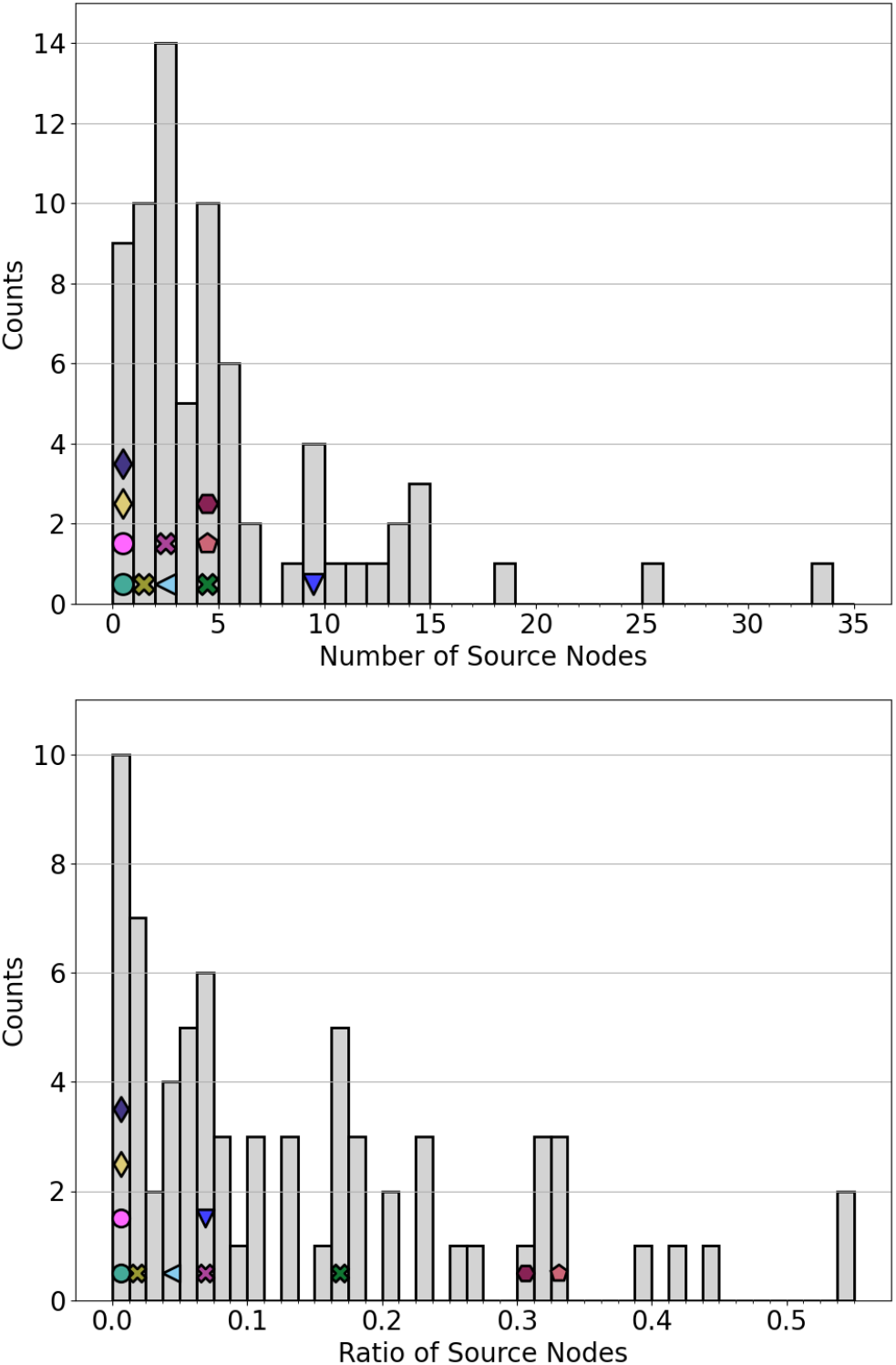
The distribution of the Cell Collective models based on the number of source nodes (top) and the ratio of source nodes (bottom).

## Appendix F: Supplementary figures

**FIG. F.1.**
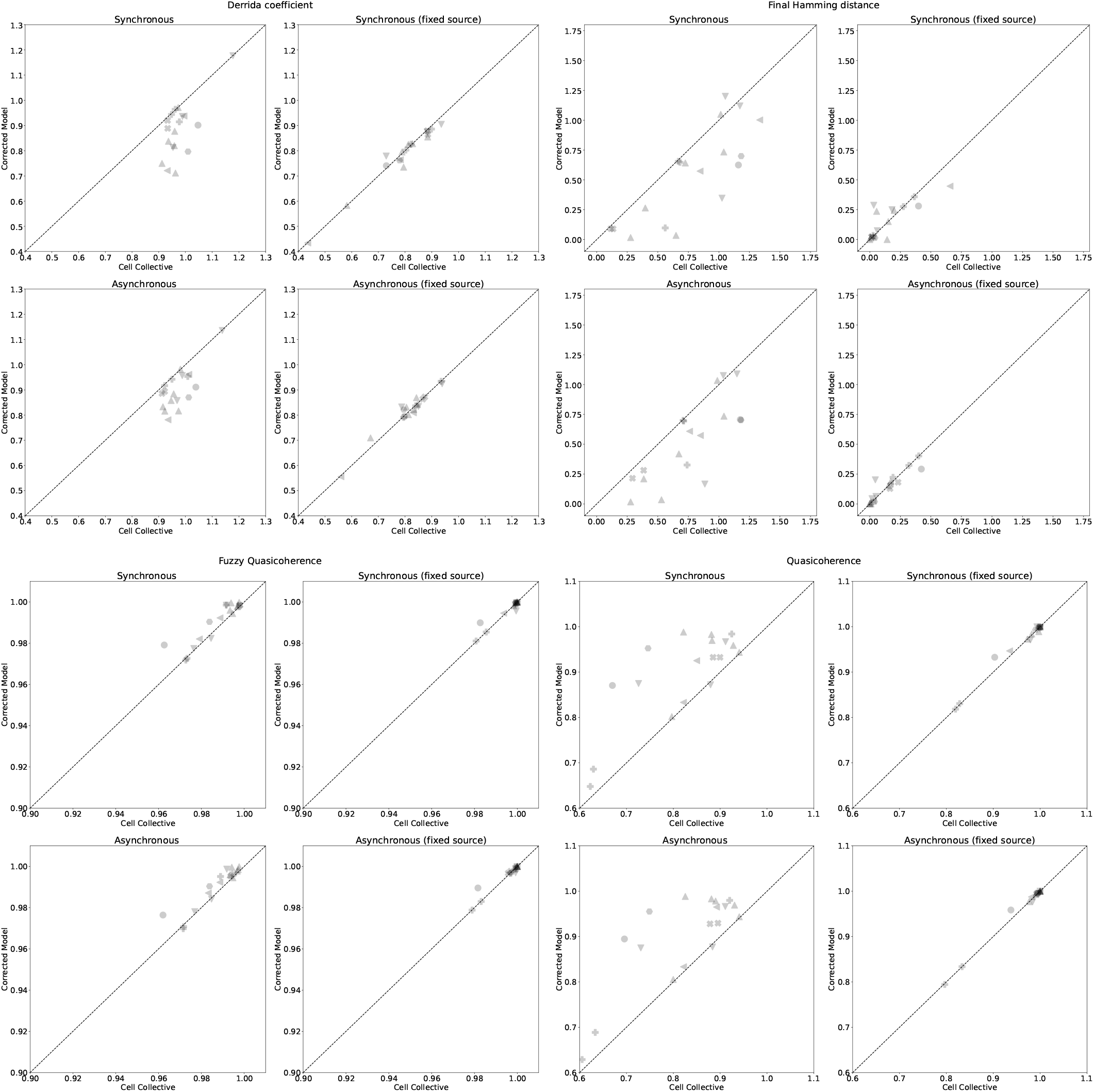
Comparison of key measures for the 18 models in the Cell Collective that were altered to attain a better agreement with the originally published models.

**FIG. F.2.**
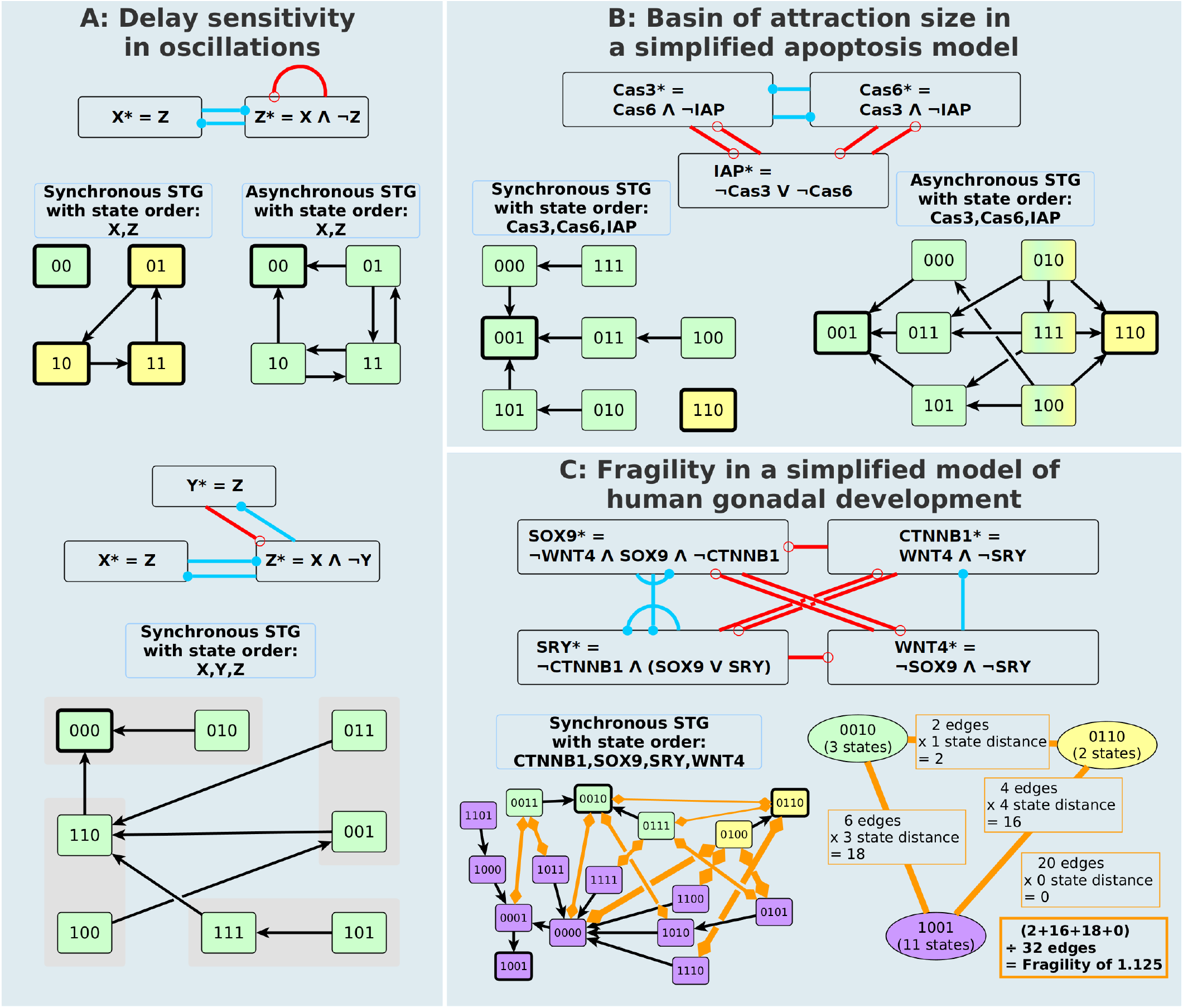
Example networks and their state transition graphs (STGs) that illustrate key points. In the interaction network, each node symbol contains the update function of the node. Blue edges ending in filled circles indicate positive regulation and red edges ending in open circles denote negative regulation. In the STGs, the attractor states are indicated by thick borders. The basin of attraction of each attractor is highlighted by the same color as the attractor. In asynchronous update, states can reach more than one attractor; such states are shaded using a gradient. In panel C the orange edges with diamond endings on each side indicate pairs of states with Hamming distance 1 that are in the basin of different attractors; these are the transitions that can arise from single-node perturbations and that lead to different long-term behavior than is observed without perturbation. The thickness of the edge indicates the Hamming distance between the corresponding attractors. The figure in the bottom right of panel C summarizes the way in which the fragility of the system can be calculated from the sum of perturbation-induced final Hamming distances between basins of attraction.

**FIG. F.3.**
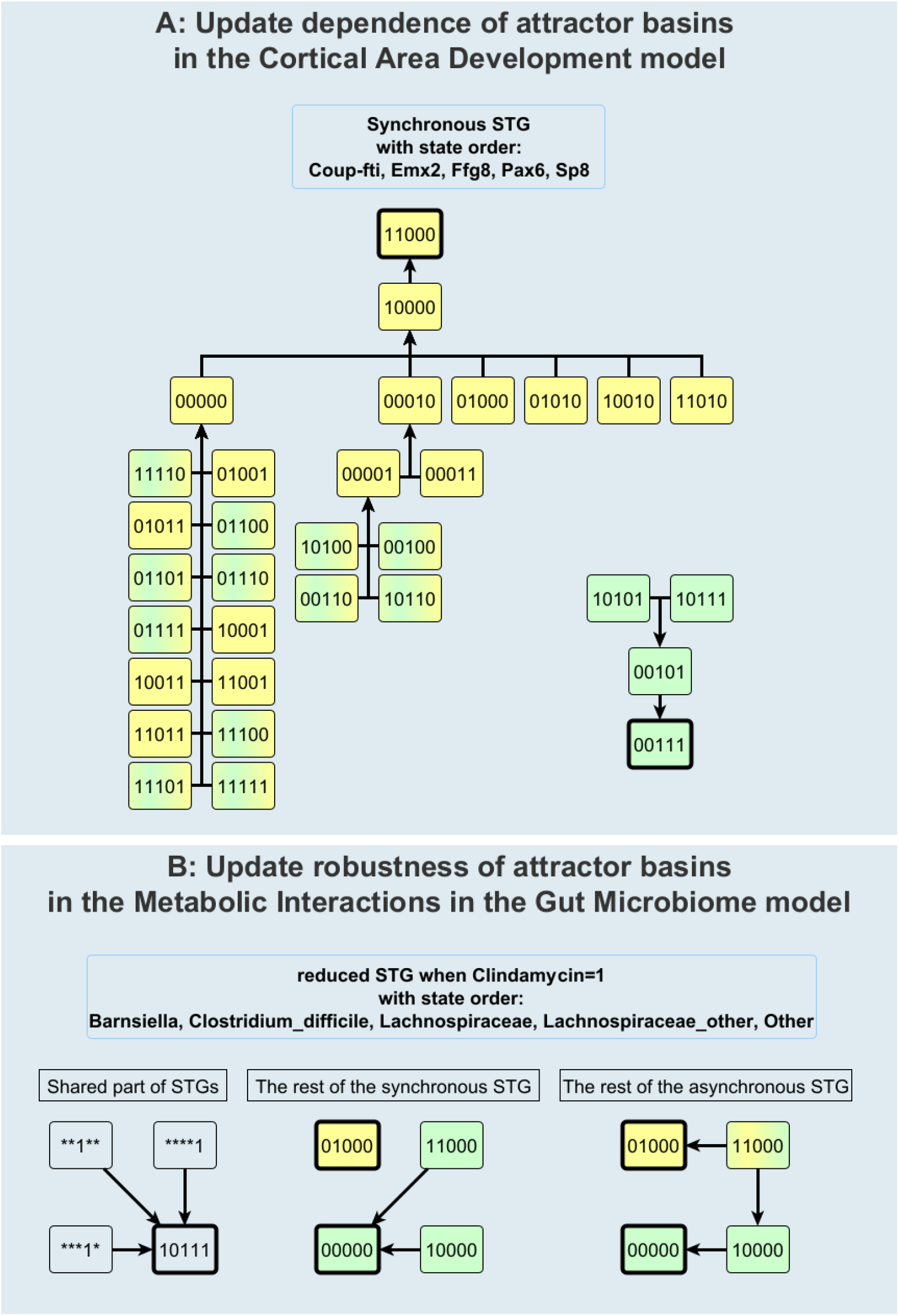
State transition graphs of (A) a model that shows a strong update dependency of attractor basins and (B) a model with update robustness of attractor basins. In both panels, node color indicates basin of attraction under asynchronous update.

**FIG. F.4.**
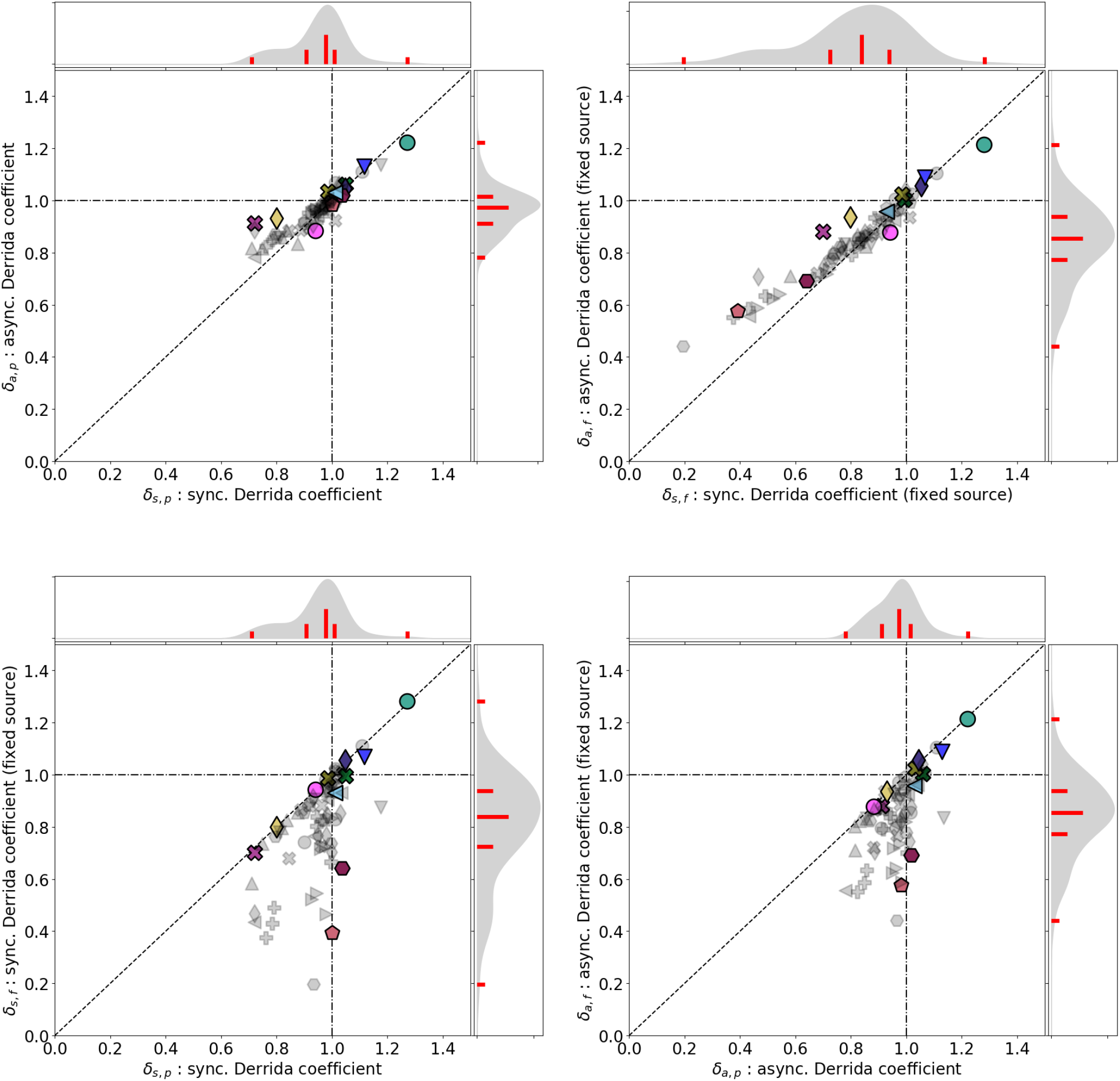
Systematic evaluation of the dependence of the Derrida coefficient *δ* on the update scheme and on source node perturbations. The ensemble of Cell Collective models shows a general agreement between the Derrida coefficients obtained for synchronous and asynchronous update (top panels). When source nodes not candidates for perturbation, the Derrida coefficient dramatically decreases (bottom panels). For example, note that three cancer drug models (plus signs) lie far from the diagonal in the lower two panels, indicating that these models are highly affected by perturbations to source nodes. This is to be expected, as the source nodes in these models represent known cancer drugs that were selected because they have a tremendous impact on the behavior of cancer cells.

**FIG. F.5.**
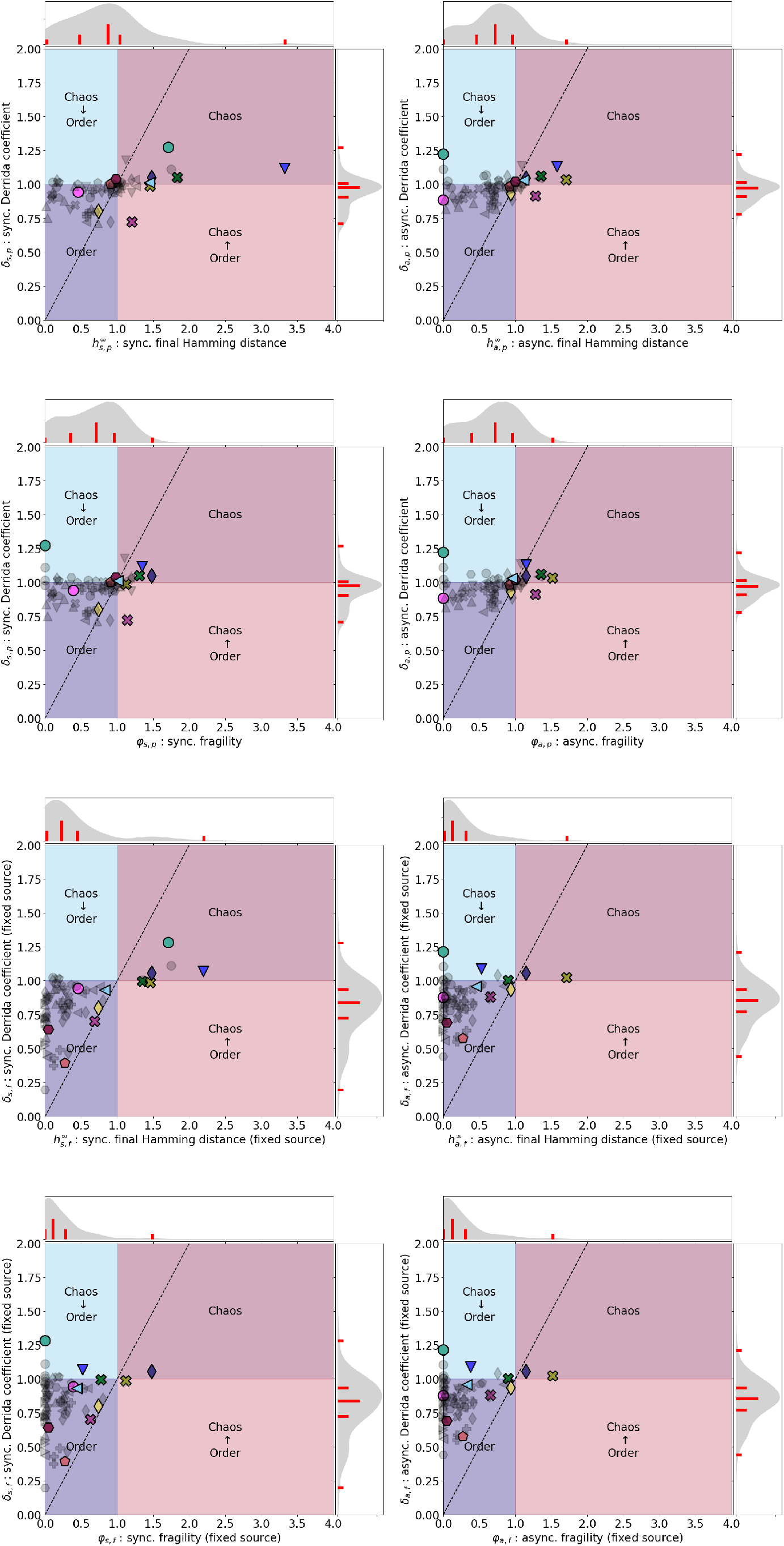
Relationships of the Derrida coefficient *δ* with the final Hamming distance *h*^∞^ and the fragility *φ*.

**FIG. F.6.**
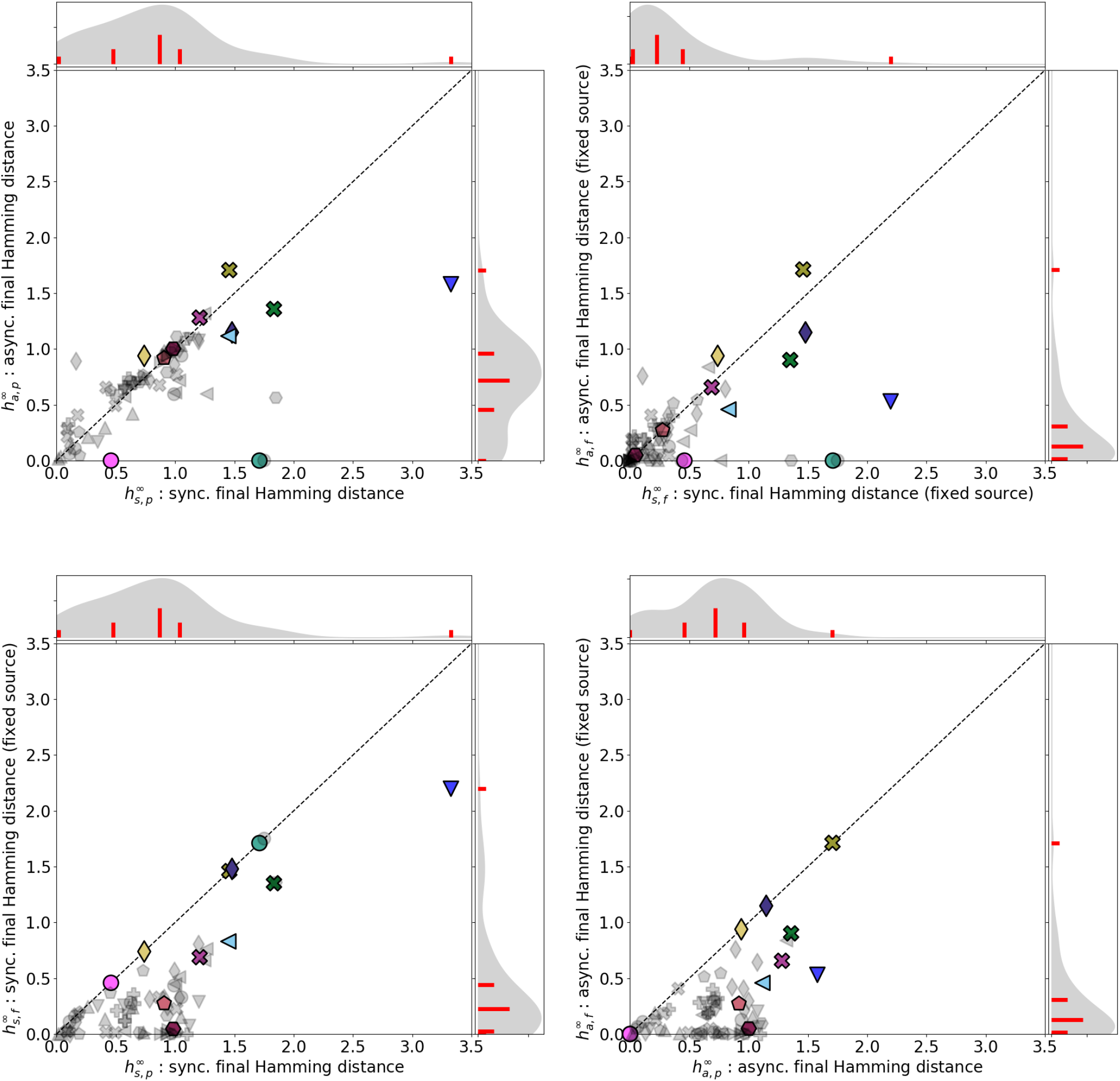
Comparison of different ways to compute *h*^∞^. The ensemble of Cell Collective models shows an overall agreement between the final Hamming distances obtained for synchronous and asynchronous update (top panels). Exceptions include models that exhibit significant phase shifts under synchronous update. When source nodes are not candidates for perturbation, the final Hamming distance dramatically decreases (bottom panels).

**FIG. F.7.**
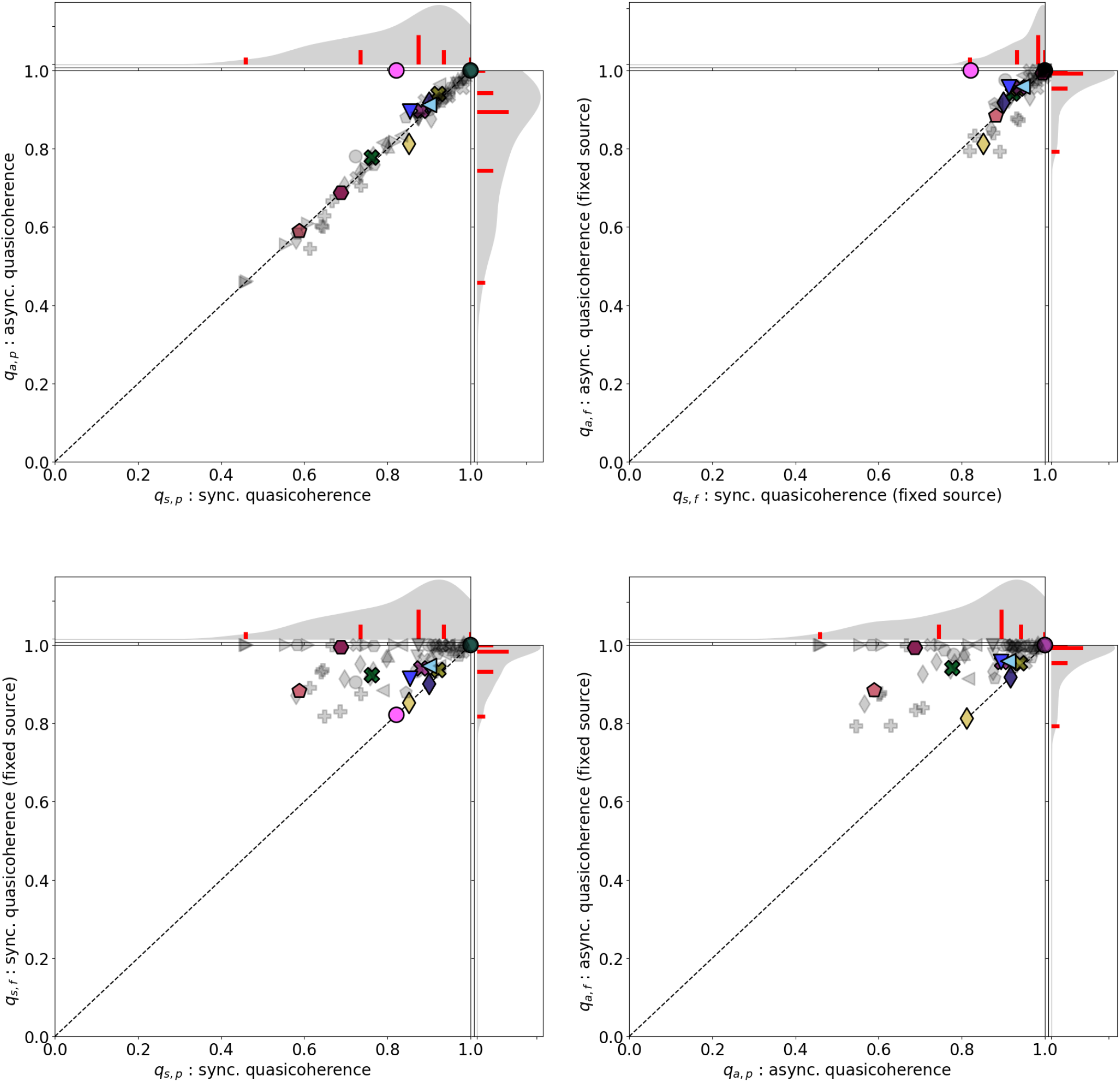
Comparison of different ways to compute *q*. There is a general agreement between the quasicoherences obtained for synchronous and asynchronous update (top panels). Making the source nodes not candidates for perturbation dramatically decreases the fragility (bottom panels).

**FIG. F.8.**
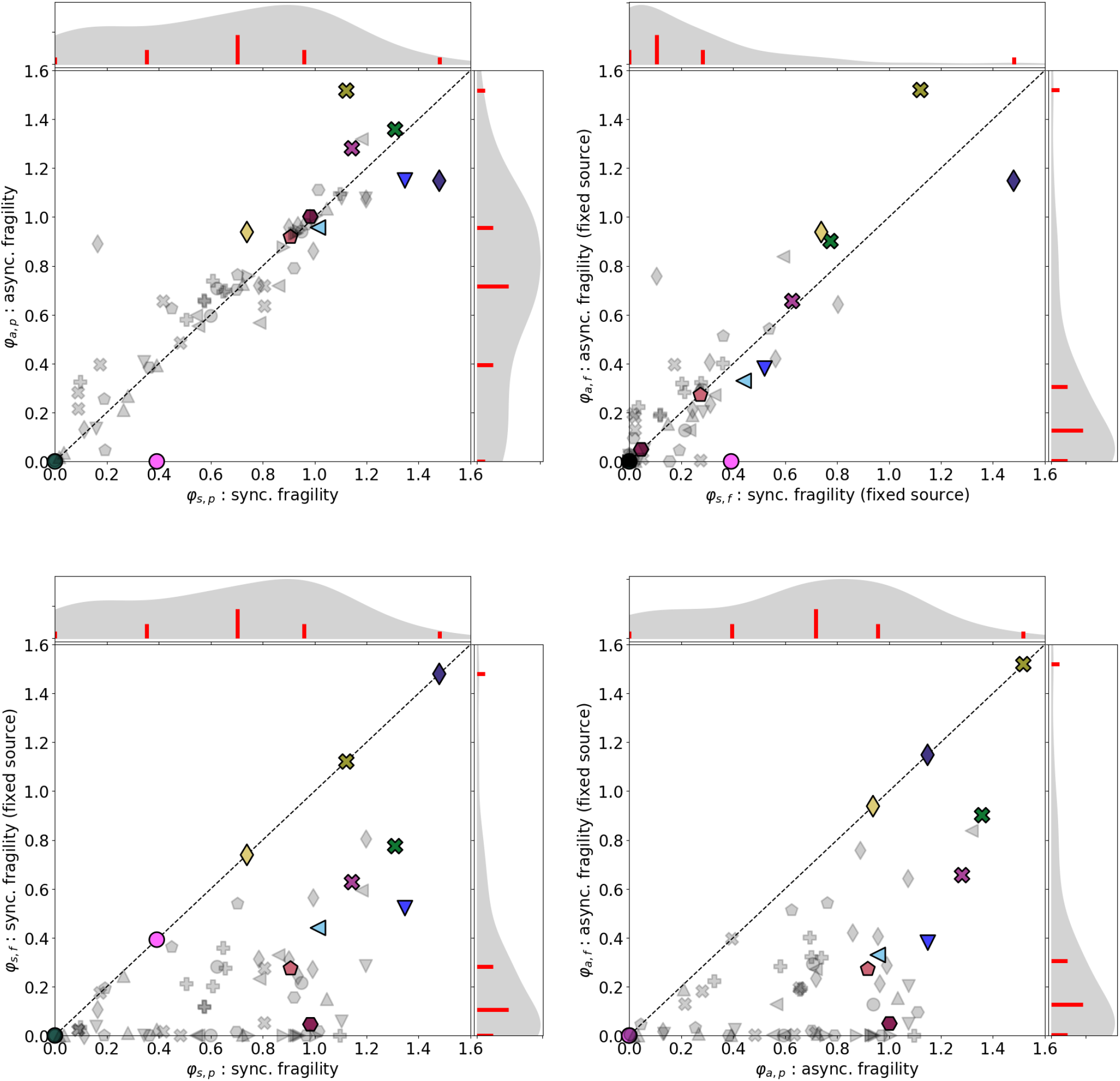
Comparison of different ways to compute *φ*. There is a general agreement between the fragilities obtained for synchronous and asynchronous update (top panels). Making the source nodes not candidates for perturbation dramatically decreases the fragility (bottom panels).

**FIG. F.9.**
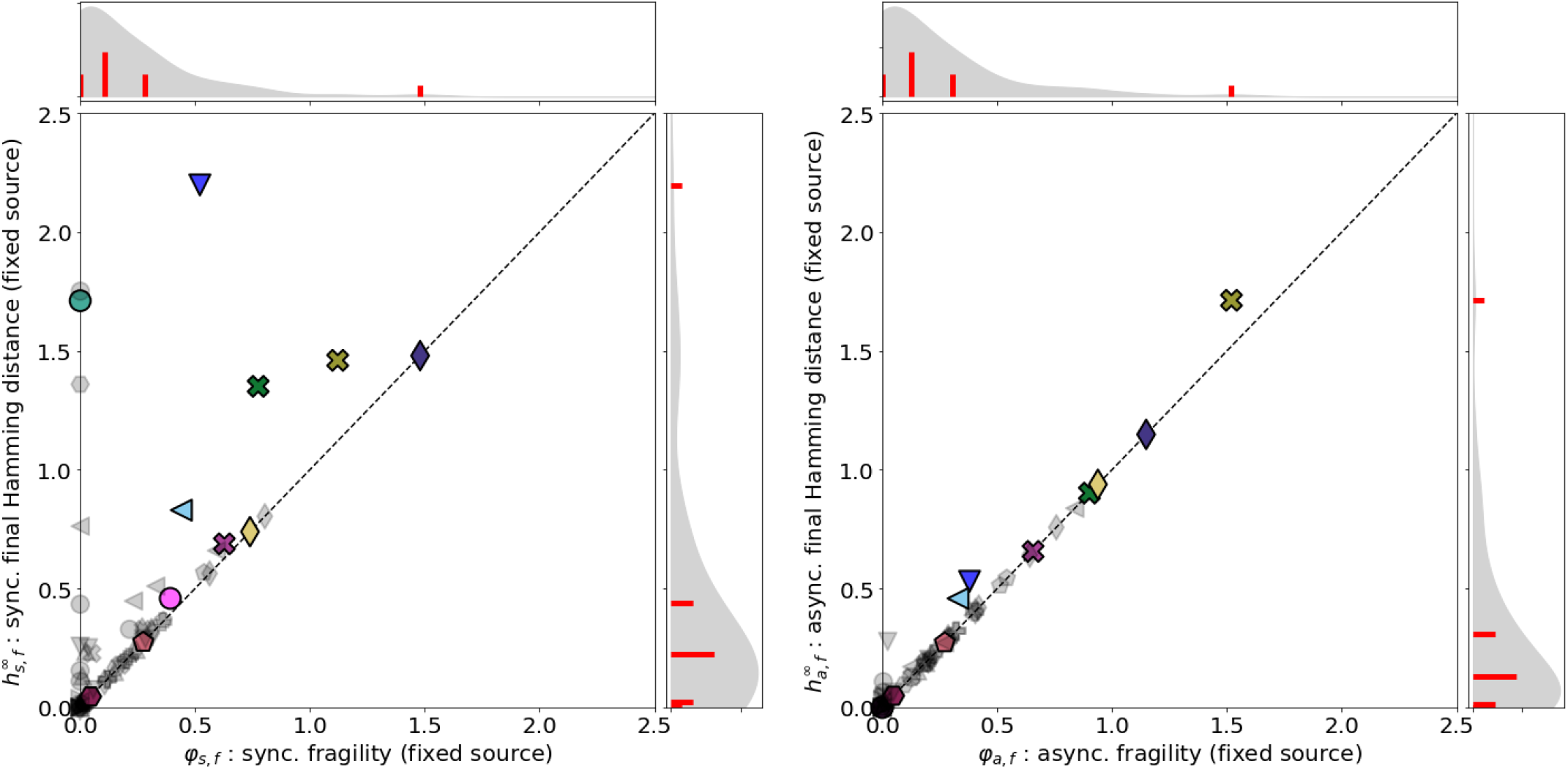
Comparison of the final Hamming distance *h*^∞^ and the fragility *φ*. The final Hamming distance is always larger than or equal to the fragility in both update schemes. The difference is much more prominent in the synchronous update in which any phase shift is permanent, compared to the asynchronous update in which the stochasticity can disperse it.

## Appendix G: Supplementary Tables

### 1. Modifications to Cell Collective models

**TABLE G.1.**
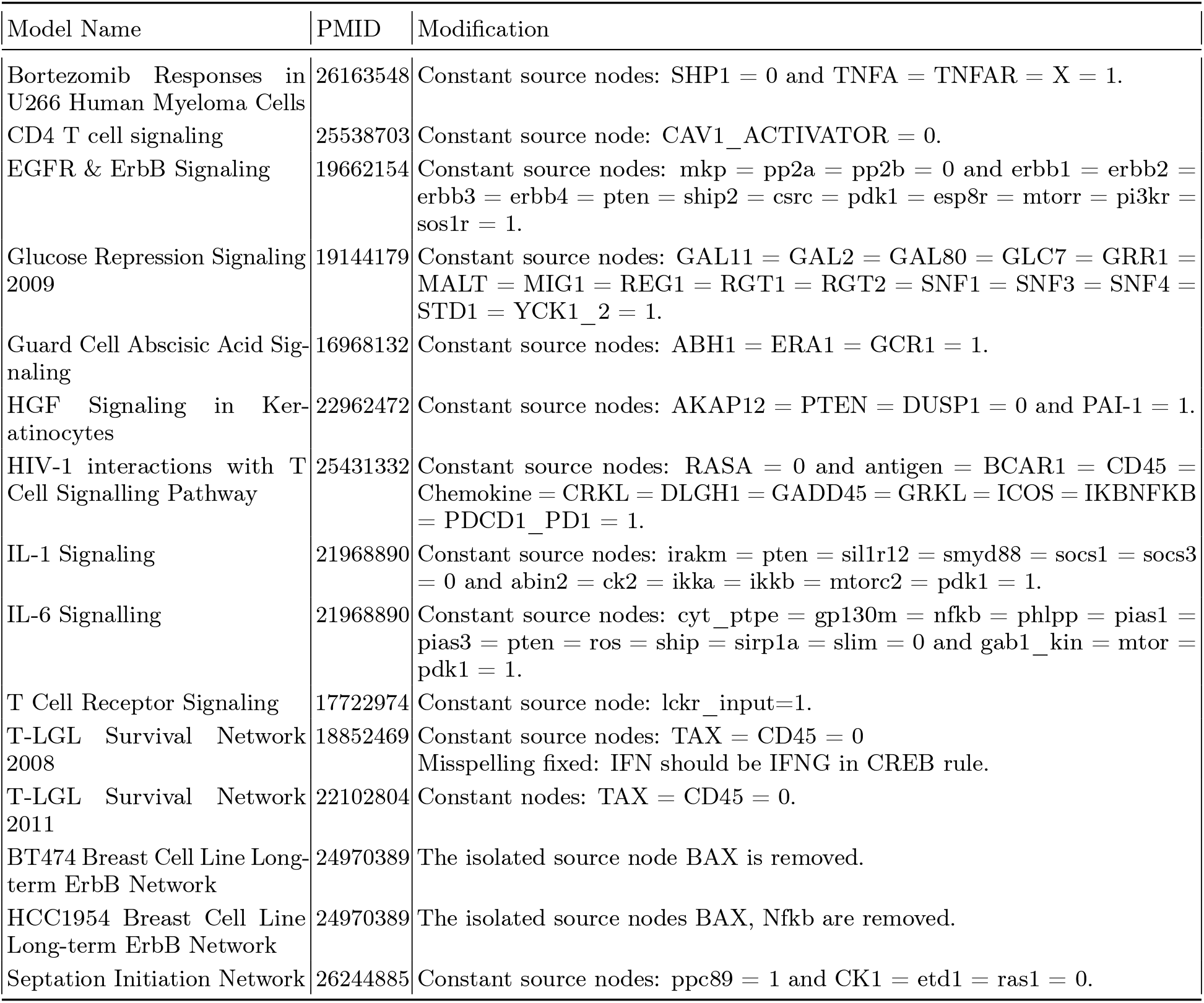
Modifications to account for source nodes that express a cellular context

**TABLE G.2.**
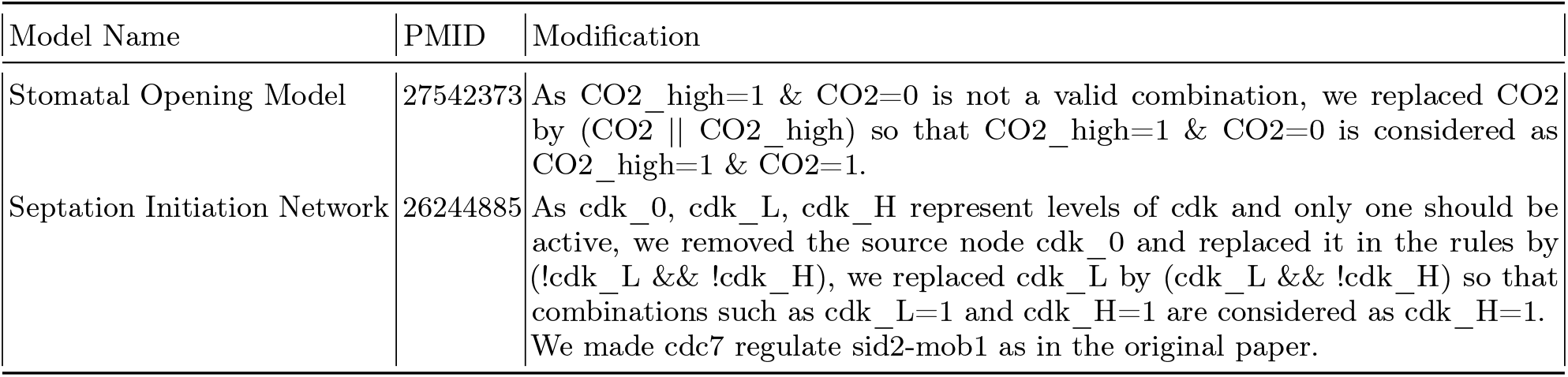
Modifications to avoid invalid combinations of source node values

**TABLE G.3.**
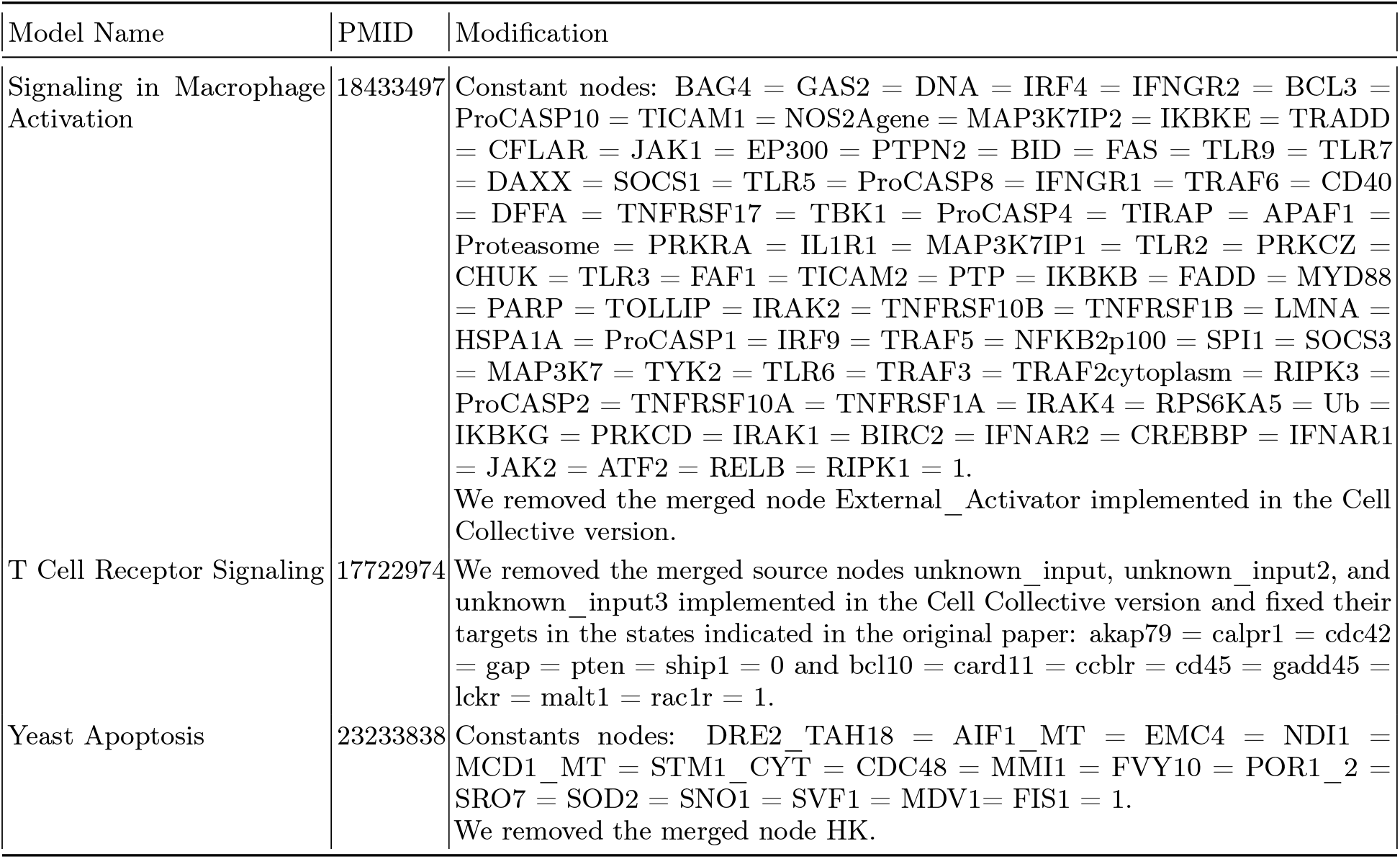
Modifications to remove aggregate source nodes and apply the original paper’s cellular context

### 2. Model characteristics

**TABLE G.4.** Cancer models

| Model Name | PMID | Nodes | Source Nodes | Mean Regulators | $\delta_{s,p}$ | $\varphi_{a,f}$ |
| --- | --- | --- | --- | --- | --- | --- |
| Aurora Kinase A in Neuroblastoma (✕) | 26616283 | 23 | 4 | 2.0435 | 1.0504 | 0.9015 |
| Colitis-associated colon cancer (✕) | 26446703 | 70 | 1 | 2.2000 | 0.9867 | 1.5197 |
| IGHV mutations in chronic lymphocytic leukemia | 26088082 | 91 | 25 | 1.3736 | 0.9615 | 0.0014 |
| Mammalian Cell Cycle | 16873462 | 20 | 1 | 2.5500 | 0.8457 | 0.1941 |
| MAPK Cancer Cell Fate Network | 24250280 | 53 | 4 | 2.0377 | 1.0051 | 0.0105 |
| Pro-inflammatory Tumor Microenvironment in Acute Lymphoblastic Leukemia | 27594840 | 26 | 2 | 3.1154 | 0.9644 | 0.0034 |
| T-LGL Survival Network 2008 | 18852469 | 60 | 4 | 3.2833 | 0.9202 | 0.1812 |
| T-LGL Survival Network 2011 Reduced Network | 22102804 | 18 | 0 | 2.3889 | 1.0125 | 0.3957 |
| T-LGL Survival Network 2011 | 22102804 | 60 | 4 | 3.3167 | 0.8886 | 0.1285 |
| Tumour Cell Invasion and Migration (✕) | 26528548 | 32 | 2 | 4.9375 | 0.7222 | 0.6563 |

**TABLE G.5.** Cancer Drug Response models

| Model Name | PMID | Nodes | Source<br>Nodes | Mean<br>Regulators | $\delta_{s,p}$ | $\varphi_{a,f}$ |
| --- | --- | --- | --- | --- | --- | --- |
| Bortezomib Responses in U266 Human Myeloma Cells | 26163548 | 67 | 1 | 1.8955 | 0.9147 | 0.2231 |
| BT474 Breast Cell Line Long-term ErbB Network | 24970389 | 24 | 5 | 3.0417 | 0.9439 | 0.3230 |
| BT474 Breast Cell Line Short-term ErbB Network | 24970389 | 16 | 5 | 3.1875 | 0.7614 | 0.1873 |
| HCC1954 Breast Cell Line Long-term ErbB Network | 24970389 | 23 | 4 | 3.1304 | 0.9643 | 0.4022 |
| HCC1954 Breast Cell Line Short-term ErbB Network | 24970389 | 16 | 5 | 3.1875 | 0.7841 | 0.1934 |
| SKBR3 Breast Cell Line Long-term ErbB Network | 24970389 | 25 | 4 | 3.4000 | 0.9458 | 0.2841 |
| SKBR3 Breast Cell Line Short-term ErbB Network | 24970389 | 16 | 5 | 2.8750 | 0.7908 | 0.3197 |
| Treatment of Castration-Resistant Prostate Cancer | 28361666 | 42 | 14 | 1.5476 | 0.9964 | 0.0000 |

**TABLE G.6.** Cell Cycle models

| Model Name | PMID | Nodes | Source<br>Nodes | Mean<br>Regulators | $\delta_{s,p}$ | $\varphi_{a,f}$ |
| --- | --- | --- | --- | --- | --- | --- |
| Arabidopsis thaliana Cell Cycle (●) | 26340681 | 14 | 0 | 4.7143 | 1.2722 | 0.0000 |
| Budding Yeast Cell Cycle 2009 | 23049686 | 18 | 0 | 3.2222 | 1.1099 | 0.0003 |
| Budding Yeast Cell Cycle | 19185585 | 20 | 4 | 2.3000 | 1.0129 | 0.1266 |
| Cell Cycle Transcription by Coupled CDK and Network Oscillators (●) | 18463633 | 9 | 0 | 2.1111 | 0.9414 | 0.0000 |
| FA BRCA pathway | 22267503 | 28 | 0 | 4.3571 | 1.0143 | 0.0036 |
| Fanconi anemia and checkpoint recovery | 26385365 | 15 | 0 | 4.2667 | 0.9783 | 0.0045 |
| Mammalian Cell Cycle 2006 | 19118495 | 10 | 1 | 3.5000 | 1.0135 | 0.0000 |
| Septation Initiation Network | 26244885 | 30 | 2 | 1.6333 | 0.9014 | 0.2923 |

**TABLE G.7.** Development and Differentiation models

| Model Name | PMID | Nodes | Source<br>Nodes | Mean<br>Regulators | $\delta_{s,p}$ | $\varphi_{a,f}$ |
| --- | --- | --- | --- | --- | --- | --- |
| B cell differentiation | 26751566 | 22 | 5 | 2.0000 | 1.0003 | 0.2118 |
| CD4+ T Cell Differentiation and Plasticity | 30116195 | 18 | 6 | 4.6667 | 0.7214 | 0.4048 |
| CD4+ T cell Differentiation | 26090929 | 38 | 9 | 2.6316 | 0.9837 | 0.2866 |
| Cortical Area Development (◆) | 20862356 | 5 | 0 | 2.8000 | 0.8011 | 0.9384 |
| Differentiation of T lymphocytes | 23743337 | 50 | 9 | 2.1200 | 0.9806 | 0.4199 |
| Human Gonadal Sex Determination (◆) | 26573569 | 19 | 0 | 3.9474 | 1.0490 | 1.1484 |
| Lymphoid and myeloid cell specification and transdifferentiation | 28584084 | 33 | 2 | 2.8788 | 1.0266 | 0.7574 |
| Lymphopoiesis Regulatory Network | 26408858 | 81 | 14 | 2.1235 | 0.9500 | 0.2332 |
| PC12 Cell Differentiation | 27148350 | 62 | 1 | 1.7581 | 0.9323 | 0.0162 |
| T cell differentiation | 6542429 | 23 | 4 | 1.6522 | 1.0314 | 0.6419 |

**TABLE G.8.** Infection and Microbiome models

| Model Name | PMID | Nodes | Source<br>Nodes | Mean<br>Regulators | $\delta_{s,p}$ | $\varphi_{a,f}$ |
| --- | --- | --- | --- | --- | --- | --- |
| *B. bronchiseptica* & *T. retortaeformis* coinfection | 2253585 | 53 | 1 | 2.5660 | 1.0004 | 0.5127 |
| *Bordetella bronchiseptica* | 2253585 | 33 | 0 | 2.3939 | 1.0137 | 0.0453 |
| Influenza A Virus Replication Cycle | 23081726 | 131 | 11 | 2.3282 | 0.9007 | 0.0313 |
| Metabolic Interactions in the Gut Microbiome (🍷) | 26102287 | 12 | 4 | 2.5833 | 1.0017 | 0.2718 |
| *Trichostrongylus retortaeformis* | 2253585 | 26 | 1 | 2.2692 | 1.0117 | 0.5418 |

**TABLE G.9.** Metabolism models

| Model Name | PMID | Nodes | Source<br>Nodes | Mean<br>Regulators | $\delta_{s,p}$ | $\varphi_{a,f}$ |
| --- | --- | --- | --- | --- | --- | --- |
| Cholesterol Regulatory Pathway | 19025648 | 34 | 2 | 1.2647 | 0.9927 | 0.0000 |
| Glucose Repression Signaling 2009 | 19144179 | 73 | 3 | 1.3699 | 0.7966 | 0.0193 |
| Iron acquisition & oxidative stress response in *A. fumigatus* | 25908096 | 22 | 2 | 1.8182 | 1.0373 | 0.0001 |
| Lac Operon | 21563979 | 13 | 3 | 1.9231 | 0.9974 | 0.0952 |
| Regulation of the L-arabinose operon of *Escherichia coli* (🍷) | 28639170 | 13 | 4 | 1.6154 | 1.0379 | 0.0492 |
| TOL Regulatory Network | 23171249 | 24 | 10 | 2.4167 | 0.9347 | 0.0000 |

**TABLE G.10.** Models of *Drosophila melanogaster* signalling pathways.

| Model Name | PMID | Nodes | Source<br>Nodes | Mean<br>Regulators | $\delta_{s,p}$ | $\varphi_{a,f}$ |
| --- | --- | --- | --- | --- | --- | --- |
| FGF pathway of *Drosophila* Signalling Pathways | 23868318 | 23 | 9 | 1.3478 | 0.9785 | 0.0000 |
| HH Pathway of *Drosophila* Signaling Pathways | 23868318 | 24 | 13 | 1.8750 | 0.9284 | 0.0000 |
| Processing of Spz Network from the *Drosophila* Signaling Pathway | 23868318 | 24 | 6 | 1.4167 | 0.9460 | 0.0000 |
| Toll Pathway of *Drosophila* Signaling Pathway | 23868318 | 11 | 2 | 1.1818 | 1.0003 | 0.0000 |
| VEGF Pathway of *Drosophila* Signaling Pathway | 23868318 | 18 | 8 | 1.4444 | 0.9604 | 0.0000 |
| Wg Pathway of *Drosophila* Signalling Pathways | 23868318 | 26 | 14 | 1.6538 | 0.9803 | 0.0019 |

**TABLE G.11.** Models of signal transduction relative to immune system cells.

| Model Name | PMID | Nodes | Source<br>Nodes | Mean<br>Regulators | $\delta_{s,p}$ | $\varphi_{a,f}$ |
| --- | --- | --- | --- | --- | --- | --- |
| CD4 T cell signaling | 25538703 | 188 | 33 | 2.0160 | 0.9713 | 0.1535 |
| HIV-1 interactions with T Cell Signalling Pathway | 25431332 | 138 | 2 | 2.2029 | 0.8771 | 0.1888 |
| IL-1 Signaling | 21968890 | 118 | 2 | 1.8644 | 0.8375 | 0.0000 |
| IL-6 Signalling | 21968890 | 86 | 1 | 1.7442 | 0.7495 | 0.0000 |
| Signaling in Macrophage Activation | 18433497 | 320 | 18 | 1.4125 | 0.7113 | 0.0066 |
| T Cell Receptor Signaling | 17722974 | 98 | 3 | 1.5102 | 0.8200 | 0.0015 |
| T-Cell Signaling 2006 | 16464248 | 40 | 3 | 1.3750 | 0.9857 | 0.0071 |

**TABLE G.12.** Models of signal transduction in stress, damage and homeostasis.

| Model Name | PMID | Nodes | Source<br>Nodes | Mean<br>Regulators | $\delta_{s,p}$ | $\varphi_{a,f}$ |
| --- | --- | --- | --- | --- | --- | --- |
| Apoptosis Network (◀) | 19422837 | 41 | 2 | 1.8293 | 1.0118 | 0.3297 |
| Death Receptor Signaling | 20221256 | 28 | 3 | 1.7143 | 1.0350 | 0.8377 |
| Guard Cell Absciscic Acid Signaling | 16968132 | 44 | 1 | 1.7955 | 0.9378 | 0.1277 |
| Oxidative Stress Pathway | 23134720 | 19 | 1 | 1.7368 | 0.9839 | 0.0001 |
| Senescence Associated Secretory Phenotype | 29206223 | 51 | 2 | 1.9216 | 0.9844 | 0.2714 |
| Yeast Apoptosis | 23233838 | 72 | 12 | 1.5278 | 0.7210 | 0.0024 |

**TABLE G.13.** Other models of signal transduction.

| Model Name | PMID | Nodes | Source<br>Nodes | Mean<br>Regulators | $\delta_{s,p}$ | $\varphi_{a,f}$ |
| --- | --- | --- | --- | --- | --- | --- |
| EGFR & ErbB Signaling | 19662154 | 104 | 13 | 2.2981 | 0.8128 | 0.0074 |
| HGF Signaling in Keratinocytes | 22962472 | 68 | 2 | 1.5441 | 0.9361 | 0.2023 |
| Neurotransmitter Signaling Pathway | 17010384 | 16 | 2 | 1.3750 | 0.9810 | 0.0234 |
| Signal Transduction in Fibroblasts (▼) | 18250321 | 139 | 9 | 3.9640 | 1.1178 | 0.3799 |
| Stomatal Opening Model | 27542373 | 49 | 5 | 3.5510 | 1.1772 | 0.0477 |

## Reference

[1] J. A and M. M. Bistability and asynchrony in a boolean model of the l-arabinose operon in escherichia coli. Bull Math Biol., 79:1778–1795, 2017

[2] W. Abou-Jaoudé, P. Traynard, P. T. Monteiro, J. Saez- Rodriguez, T. Helikar, D. Thieffry, and C. Chaouiya. Logical Modeling and Dynamical Analysis of Cellular Networks. Frontiers in Genetics, 7, 2016. ISSN 1664-8021. URL https://www.frontiersin.org/articles/10.3389/fgene.2016.00094

[3] I. Albert, J. Thakar, S. Li, R. Zhang, and R. Albert. Boolean network simulations for life scientists. Source Code Biol Med, 3(16), 2008. doi:10.1186/1751-0473-3-16.

[4] M. Aldana and P. Cluzel. A natural class of robust networks. Proc Natl Acad Sci U S A, 100(15):8710–8714, 2003. doi:110.1073/pnas.1536783100.

[5] S.-i. Azuma, T. Yoshida, and T. Sugie. Structural monos-tability of activation-inhibition boolean networks. IEEE Transactions on Control of Network Systems, 4(2):179–190, 2017. doi:10.1109/TCNS.2015.2485440.

[6] E. Balleza, E. R. Alvarez-Buylla, A. Chaos, S. Kauffman, I. Shmulevich, and M. Aldana. Critical Dynamics in Genetic Regulatory Networks: Examples from Four Kingdoms. PLOS ONE, 3(6):e2456, June 2008. ISSN 1932-6203. doi:10.1371/journal.pone.0002456. URL https://journals.plos.org/plosone/article?id=10.1371/journal.pone.0002456. Publisher: Public Library of Science

[7] I. Barbaric, G. Miller, and T. N. Dear. Appearances can be deceiving: phenotypes of knockout mice. Briefings in Functional Genomics, 6(2):91–103, 06 2007. ISSN 2041-2649. doi:10.1093/bfgp/elm008. URL https://doi.org/10.1093/bfgp/elm008

[8] D. P. A. Cohen, L. Martignetti, S. Robine, E. Barillot, A. Zinovyev, and L. Calzone. Mathematical modelling of molecular pathways enabling tumour cell invasion and migration. PLOS Computational Biology, 11(11):1–29, 11 2015. doi:10.1371/journal.pcbi.1004571. URL https://doi.org/10.1371/journal.pcbi.1004571

[9] S. Collombet, C. van Oevelen, J. L. Sardina Ortega, W. Abou-Jaoudé, B. Di Stefano, M. Thomas-Chollier, T. Graf, and D. Thieffry. Logical modeling of lymphoid and myeloid cell specification and trans-differentiation. Proceedings of the National Academy of Sciences, 114(23):5792–5799, June 2017. doi: 10.1073/pnas.1610622114. URL https://www.pnas.org/doi/full/10.1073/pnas.1610622114. Publisher: Proceedings of the National Academy of Sciences

[10] M. Conrad. Adaptability: The significance of variability from molecule to ecosystem. Springer Science & Business Media, 2012

[11] R. B. Correia, A. J. Gates, X. Wang, and L. M. Rocha. Cana: A python package for quantifying control and canalization in boolean networks. Frontiers in Physiology, 9, 2018. doi:10.3389/fphys.2018.01046.

[12] F. X. Costa, J. C. Rozum, A. M. Marcus, and L. M. Rocha. Effective connectivity and bias entropy improve prediction of dynamical regime in automata networks. Entropy, 25 (2), 2023. ISSN 1099-4300. doi:10.3390/e25020374. URL https://www.mdpi.com/1099-4300/25/2/374

[13] M. Dahlhaus, A. Burkovski, F. Hertwig, C. Mussel, R. Volland, M. Fischer, K.-M. Debatin, H. A. Kestler, and C. Beltinger. Boolean modeling identifies greatwall/mastl as an important regulator in the aurka network of neuroblastoma. Cancer Letters, 371(1):79–89, 2016. ISSN 0304-3835. doi: https://doi.org/10.1016/j.canlet.2015.11.025. URL https://www.sciencedirect.com/science/article/pii/S0304383515007016

[14] B. C. Daniels, H. Kim, D. Moore, S. Zhou, H. B. Smith, B. Karas, S. A. Kauffman, and S. I. Walker. Criticality distinguishes the ensemble of biological regulatory networks. Phys Rev Lett., 121(138102), 2005. doi: 10.1103/PhysRevLett.121.138102.

[15] D. Deritei, J. Rozum, E. Ravasz Regan, and R. Albert. A feedback loop of conditionally stable circuits drives the cell cycle from checkpoint to checkpoint. Scientific Reports, 9(16430), 2019

[16] B. Derrida and Y. Pomeau. Random networks of automata: a simple annealed approximation. EPL (Euro-physics Letters), 1(2):45, 1986

[17] B. Derrida and D. Stauffer. Phase transitions in two-dimensional kauffman cellular automata. EPL (Euro-physics Letters), 2(10):739, 1986

[18] B. Drossel. Random boolean networks. In H. G. Schuster, editor, Reviews of Nonlinear Dynamics and Complexity, pages 69–110. Wiley, 2008

[19] A. Fauré, A. Naldi, C. Chaouiya, and D. Thieffry. Dynamical analysis of a generic boolean model for the control of the mammalian cell cycle. Bioinformatics, 22:e124–131, 2006. doi:10.1093/bioinformatics/btl210.

[20] X. Gan and R. Albert. Analysis of a dynamic model of guard cell signaling reveals the stability of signal propagation. BMC systems biology, 10:1–14, 2016

[21] A. Garg, A. Di Cara, I. Xenarios, L. Mendoza, and G. De Micheli. Synchronous versus asynchronous modeling of gene regulatory networks. Bioinformatics, 24(17):1917–1925, 07 2008. ISSN 1367-4803. doi: 10.1093/bioinformatics/btn336. URL https://doi.org/10.1093/bioinformatics/btn336

[22] A. J. Gates, R. Brattig Correia, X. Wang, and L. M. Rocha. The effective graph reveals redundancy, canalization, and control pathways in biochemical regulation and signaling. Proceedings of the National Academy of Sciences, 118(12):e2022598118, 2021

[23] C. Gershenson. Updating schemes in random boolean networks. In Artificial Life IX: Proceedings of the Ninth International Conference on the Simulation and Synthesis of Living Systems, pages 238–243. MIT Press, 2004

[24] C. E. Giacomantonio and G. J. Goodhill. A boolean model of the gene regulatory network underlying mammalian cortical area development. PLOS Computational Biology, 6 (9):1–13, 09 2010. doi:10.1371/journal.pcbi.1000936. URL https://doi.org/10.1371/journal.pcbi.1000936

[25] G. Giaever, A. Chu, and L. e. a. Ni. Functional profiling of the saccharomyces cerevisiae genome. Nature, 418: 387—-391, 2006. doi:10.1038/nature00935.

[26] F. Greil and B. Drossel. Dynamics of critical kauffman networks under asynchronous stochastic update. Phys. Rev. Lett., 95:048701, Jul 2005. doi: 10.1103/PhysRevLett.95.048701. URL https://link.aps.org/doi/10.1103/PhysRevLett.95.048701

[27] S. Gupta, S. S. Bisht, R. Kukreti, S. Jain, and S. K. Brahmachari. Boolean network analysis of a neurotransmitter signaling pathway. Journal of Theo-retical Biology, 244(3):463–469, 2007. ISSN 0022-5193. doi:https://doi.org/10.1016/j.jtbi.2006.08.014. URL https://www.sciencedirect.com/science/article/pii/S0022519306003675

[28] T. Helikar, J. Konvalina, J. Heidel, and J. A. Rogers. Emergent decision-making in biological signal transduction networks. Proc Natl Acad Sci U S A, 105:1913–1918, 2008. doi:10.1073/pnas.0705088105.

[29] T. Helikar, B. Kowal, S. McClenathan, M. Bruckner, T. Rowley, A. Madrahimov, B. Wicks, M. Shrestha, K. Limbu, and J. A. Rogers. The cell collective: toward an open and collaborative approach to systems biology. BMC systems biology, 6(1):1–14, 2012

[30] C. Kadelka, T.-M. Butrie, E. Hilton, J. Kinseth, and H. Serdarevic. A meta-analysis of Boolean network models reveals design principles of gene regulatory networks, Sept. 2020. URL http://arxiv.org/abs/2009.01216. arXiv:2009.01216 [nlin, q-bio]

[31] R. Kamath, A. Fraser, Y. Dong, G. Poulin, R. Durbin, M. Gotta, A. Kanapin, N. Le Bot, S. Moreno, M. Sohrmann, D. P. Welchman, P. Zipperlen, and J. Ahringer. Systematic functional analysis of the caenorhabditis elegans genome using rnai. Nature, 421: 231–237, 2003. doi:10.1038/nature01278.

[32] S. Kauffman. Gene regulation networks: A theory for their global structure and behaviors. Current Topics in Developmental Biology, 6, 1971

[33] H. Klarner, A. Streck, and H. Siebert. PyBool-Net: a python package for the generation, analysis and visualization of boolean networks. Bioinformatics, 33(5):770–772, 12 2016. ISSN 1367-4803. doi: 10.1093/bioinformatics/btw682. URL https://doi.org/10.1093/bioinformatics/btw682

[34] S. Li, S. M. Assmann, and R. Albert. Predicting essential components of signal transduction networks: A dynamic model of guard cell abscisic acid signaling. PLOS Biology, 4(10):1–17, 09 2006. doi: 10.1371/journal.pbio.0040312. URL https://doi.org/10.1371/journal.pbio.0040312

[35] J. Lu, H. Zeng, Z. Liang, L. Chen, L. Zhang, H. Zhang, H. Liu, H. Jiang, B. Shen, M. Huang, M. Geng, S. Spiegel, and C. Luo. Network modelling reveals the mechanism underlying colitis-associated colon cancer and identifies novel combinatorial anti-cancer targets. Sci Rep., 8:14739, 2015. doi:10.1038/srep14739.

[36] Z. Mai and H. Liu. Boolean network-based analysis of the apoptosis network: Irreversible apoptosis and stable surviving. Journal of Theoretical Biology, 259(4):760–769, 2009. ISSN 0022-5193. doi:https://doi.org/10.1016/j.jtbi.2009.04.024. URL https://www.sciencedirect.com/science/article/pii/S0022519309001878

[37] S. Manicka, M. Marques-Pita, and L. Rocha. Effective connectivity determines the critical dynamics of biochemical networks. J. R. Soc. Interface, 19(20210659), 2022. doi:10.1098/rsif.2021.0659.

[38] A. Mbodj, G. Junion, C. Brun, E. Furlong, and D. Thieffry. Logical modelling of drosophila signalling pathways. Mol Biosyst., 9:2248–2258, 2013. doi:10.1039/c3mb70187e.

[39] A. A. Moreira and L. A. N. Amaral. Canalizing Kauffman Networks: Nonergodicity and Its Effect on Their Critical Behavior. Physical Review Letters, 94(21): 218702, June 2005. ISSN 0031-9007, 1079–7114. doi: 10.1103/PhysRevLett.94.218702. URL https://link.aps.org/doi/10.1103/PhysRevLett.94.218702

[40] A. Naldi, E. Remy, D. Thieffry, and C. Chaouiya. Dynamically consistent reduction of logical regulatory graphs. Theoretical Computer Science, 412(21):2207–2218, 2011. ISSN 0304-3975. doi: https://doi.org/10.1016/j.tcs.2010.10.021. URL https://www.sciencedirect.com/science/article/pii/S0304397510005839. Selected Papers from the 7th International Conference on Computational Methods in Systems Biology

[41] R. Okuta, Y. Unno, D. Nishino, S. Hido, and C. Loomis. Cupy: A numpy-compatible library for nvidia gpu calculations. In Proceedings of Workshop on Machine Learning Systems (LearningSys) in The Thirty-first Annual Conference on Neural Information Processing Systems (NIPS), 2017. URL http://learningsys.org/nips17/assets/papers/paper_16.pdf

[42] D. A. Orlando, C. Y. Lin, A. Bernard, J. Y. Wang, J. E. Socolar, E. S. Iversen, A. J. Hartemink, and H. S. B. Global control of cell-cycle transcription by coupled cdk and network oscillators. Nature, 453:944–947, 2008. doi: 10.1038/nature06955.

[43] E. Ortiz-Gutiérrez, K. García-Cruz, E. Azpeitia, A. Castillo, M. d. l. P. Sánchez, and E. R. Álvarez-Buylla. A dynamic gene regulatory network model that recovers the cyclic behavior of arabidopsis thaliana cell cycle. PLOS Computational Biology, 11(9):1–28, 09 2015. doi:10.1371/journal.pcbi.1004486. URL https://doi.org/10.1371/journal.pcbi.1004486

[44] N. H. Packard. Adaptation toward the edge of chaos. In A. J. Mandell, M. F. Shlesinger, and J. A. S. Kelso, editors, Dynamic Patterns In Complex Systems-Proceedings Of The Conference In Honor Of Hermann Haken’s 60th Birthday, pages 293–301. World Scientific, April 1988

[45] L. Paulevé, J. Kolcák, T. Chatain, and S. Haar. Reconciling qualitative, abstract, and scalable modeling of biological networks. Nat Commun, 11(4256), 2020. doi: 10.1038/s41467-020-18112-5.

[46] O. Ríos, S. Frias, A. Rodríguez, S. Kofman, H. Merchant, L. Torres, and L. Mendoza. A boolean network model of human gonadal sex determination. Theor Biol Med Model, 12:26, 2015. doi:10.1186/s12976-015-0023-0.

[47] J. C. Rozum, J. G. T. Zanudo, X. Gan, D. Deritei, and R. Albert. Parity and time reversal elucidate both decision-making in empirical models and attractor scaling in critical boolean networks. Science Advances, 7(29):eabf8124, 2021. doi: 10.1126/sciadv.abf8124. URL https://www.science.org/doi/abs/10.1126/sciadv.abf8124

[48] K. S. Metabolic stability and and epigenesis in randomly constructed genetic nets. J Theor Biol., 22:437–467, 1969

[49] A. Saadatpour, I. Albert, and R. Albert. Attractor analysis of asynchronous boolean models of signal transduction networks. Journal of Theoretical Biology, 266(4):641–656, 2010. ISSN 0022-5193. doi:https://doi.org/10.1016/j.jtbi.2010.07.022. URL https://www.sciencedirect.com/science/article/pii/S0022519310003796

[50] M. Sales-Pardo, R. Guimera, A. A. Moreira, and L. A. N. Amaral. Extracting the hierarchical organization of complex systems. Proceedings of the National Academy of Sciences, 104(39):15224–15229, 2007

[51] I. Shmulevich and S. A. Kauffman. Activities and sensitivities in boolean network models. Phys Rev Lett., 93 (048701), 2004. doi:10.1103/PhysRevLett.93.048701.

[52] I. Shmulevich, S. A. Kauffman, and M. Aldana. Eukaryotic cells are dynamically ordered or critical but not chaotic. Proceedings of the National Academy of Sciences USA, 102:13439–13444, 2005. doi: 10.1073/pnas.0506771102

[53] S. N. Steinway, M. B. Biggs, T. P. Loughran, Jr, J. A. Papin, and R. Albert. Inference of network dynamics and metabolic interactions in the gut microbiome. PLOS Computational Biology, 11(6):1–25, 06 2015. doi: 10.1371/journal.pcbi.1004338. URL https://doi.org/10.1371/journal.pcbi.1004338

[54] G. Stoll, B. Caron, E. Viara, A. Dugourd, A. Zinovyev, A. Naldi, G. Kroemer, E. Barillot, and L. Calzone. Ma-BoSS 2.0: an environment for stochastic Boolean modeling. Bioinformatics, 33(14):2226–2228, 03 2017. ISSN 1367-4803. doi:10.1093/bioinformatics/btx123. URL https://doi.org/10.1093/bioinformatics/btx123

[55] A. Subbaroyan, O. C. Martin, and A. Samal. Minimum complexity drives regulatory logic in Boolean models of living systems. PNAS Nexus, 1(1), 04 2022. ISSN 2752-6542. doi:10.1093/pnasnexus/pgac017. URL https://doi.org/10.1093/pnasnexus/pgac017.pgac017

[56] C. Teuscher. Revisiting the edge of chaos: Again? Biosystems, 218:104693, 2022. ISSN 0303-2647. doi: https://doi.org/10.1016/j.biosystems.2022.104693. URL https://www.sciencedirect.com/science/article/pii/S0303264722000806

[57] A. Veliz-Cuba. Reduction of boolean network models. Journal of Theoretical Biology, 289:167–172, 2011. ISSN 0022-5193. doi:https://doi.org/10.1016/j.jtbi.2011.08.042. URL https://www.sciencedirect.com/science/article/pii/S0022519311004450

[58] C. H. Waddington. Canalization of development and the inheritance of acquired characters. Nature, 150(3811): 563–565, 1942

[59] K. Willadsen, J. Triesch, and J. Wiles. Understanding robustness in random boolean networks. In Artificial Life XI: Proceedings of the Eleventh International Conference on the Simulation and Synthesis of Living Systems, pages 695–701. MIT Press, 2008

[60] J. G. T. Zanudo and R. Albert. An effective network reduction approach to find the dynamical repertoire of discrete dynamic networks. Chaos: An Interdisciplinary Journal of Nonlinear Science, 23(2), 06 2013. ISSN 1054-1500. doi:10.1063/1.4809777. URL https://doi.org/10.1063/1.4809777.025111

[61] J. Zañudo, M. Aldana, and G. Martínez-Mekler. Boolean threshold networks: Virtues and limitations for biological modeling. In S. Niiranen and A. Ribeiro, editors, Information Processing and Biological Systems, volume 11 of Intelligent Systems Reference Library. Springer, 2011. doi:10.1007/978-3-642-19621-8_6

